# Predicting animal movement with deepSSF: a deep learning step selection framework

**DOI:** 10.1101/2025.02.13.638055

**Authors:** Scott W. Forrest, Dan Pagendam, Conor Hassan, Christopher Drovandi, Jonathan R. Potts, Michael Bode, Andrew J. Hoskins

## Abstract

1. Predictions of animal movement are vital for understanding and managing wild populations. However, the fine-scale, complex decision-making of animals can pose challenges for the accurate prediction of trajectories. Step selection functions (SSFs), a common tool for inferring relationships between animal movement and the environment, are also increasingly used to simulate animal trajectories for prediction. Although admitting a lot of flexibility, the SSF framework is limited to its reliance on pre-defined functional forms for fitting to data, and SSFs that involve complex functional forms to model detailed processes can be prohibitively difficult to fit and interpret.
2. Here, we present `deepSSF`, an approach to fit and predict animal movement data using deep learning. Whilst not specific to any particular model, we denote the `deepSSF` approach as building and training a neural network architecture that receives multiple environmental layers and scalar values as inputs, and outputs a single layer representing the next-step probability. To demonstrate a `deepSSF` model, we build a model in `PyTorch` that has distinct but interacting habitat selection and movement subnetworks, which allows for explicit representation of both processes and interpretable intermediate outputs. We apply our model to GPS data of introduced water buffalo (*Bubalus bubalis*) in the tropical savannas of Northern Australia.
3. Our `deepSSF` model was able to learn features that are present in the habitat covariate layers, such as linear features (rivers, forest edges), and the composition of certain habitat areas, without having to specify them pre-emptively within the SSF framework. It was also able to capture the complex interactions between the habitat covariates as well as temporal dynamics across time of day and year.
4. We expect that the `deepSSF` approach will generate accurate and informative animal movement trajectories, which can be used for deepening our understanding of animal-environment systems and for the practical management of species. Considering the wide range of deep learning tools, the `deepSSF` approach could be extended to represent memory and social dynamic processes, with the potential for integrating non-spatial data sources such as accelerometers and physiological sensors.

## 1 Introduction

The movement of animals through space drives the structure and functioning of ecosystems through processes such as seed dispersal, herbivory and predator-prey relationships (Fortin et al., 2005; Côrtes & Uriarte, 2013; Bello et al., 2024). An ability to predict animal locations is also vital for conservation efforts. Accurate predictions of movement paths and locations can be used to mitigate impacts to wildlife when designing infrastructure such as roads, wind turbines or mines (Mayer et al., 2021; Cowan et al., 2024), to reduce the risk to threatened species (Finnegan et al., 2021; Forrest, Rodríguez-Recio, & Seddon, 2024), or to manage invasive or problematic species (Lustig et al., 2019; Pili et al., 2022; Patterson et al., 2024).

Recently, there has been a growing prevalence of predicting animal movement by simulating it from fitted step selection functions (SSFs), a flexible way of incorporating measured processes such as movement, habitat selection, memory and social interactions (Signer et al., 2017; Potts & Börger, 2023; Signer et al., 2023; Ellison, Potts, Boudreau, et al., 2024; Ellison, Potts, Strickland, et al., 2024). SSFs have been used in this context for quantifying connectivity (Hooker et al., 2021; Whittington et al., 2022; Hofmann et al., 2023; Sells et al., 2023), expected distributions (Forrest, Pagendam, et al., 2024), and the effect of different landscapes on movement behaviour (MA Cowan, unpublished data). Simulating from an SSF requires proposing potential steps from a starting location, determining the relative probability of choosing each of these steps (using the fitted movement and habitat selection parameters), and then choosing one of them using these relative probabilities. The end location of this step then becomes the next step’s starting point and the process is repeated. Recent developments aim to increase the flexibility of SSFs through the addition of temporally dynamic and flexible responses (Forrest, Pagendam, et al., 2024; Klappstein et al., 2024), which can increase the realism of the trajectories. However, animal movement is notoriously difficult to predict at fine scales, and the decisions that animals make depend on a range of factors which may interact in ways that are difficult to predict *a priori* (Nathan et al., 2008; Couzin, 2009; Strandburg-Peshkin et al., 2015). Therefore, to make accurate predictions, we require a framework that can handle complex interactions and does not require pre-emptive specification of functional forms and responses for animal movement. One such approach is deep learning, a highly flexible data-driven method that can accommodate a large number of inputs and represent an impressively wide range of complex processes (LeCun et al., 2015; Drori, 2022).

Deep learning uses multiple processing layers to construct an internal representation of data that can have multiple layers of abstraction (LeCun et al., 2015). It has led to many advances in both our understanding of the world around us, and our ability to generate predictions of complex and abstract phenomena (Raghu & Schmidt, 2020). In recent years deep learning has advanced rapidly, with progress in both modelling architectures that enable more flexible model fitting, but also hardware such as Graphics Processing Units (graphics processing units) that have enabled models to be fitted with much greater speed. Increasingly, deep learning is being applied to complex problems in science, often with impressive results (Raghu & Schmidt, 2020; Jumper et al., 2021). The flexibility of deep learning models, and their ability to take an array of different inputs and produce almost any output, means that with some thought and consideration they can help uncover interesting features in almost any dataset. There are many existing applications of deep learning that are similar to the next-step-ahead predictions of animal movement, such as the movement of a robot, game playing, text generation and the generation of human and vehicle trajectories (Drori, 2022). Although the application of deep learning has yet to be fully utilised or adopted for modelling animal movement, there have been some promising examples that consider next-step ahead predictions using a range of machine learning architectures (Dalziel et al., 2008; Wijeyakulasuriya et al., 2020; Shenk et al., 2021; Cífka et al., 2023; Chen et al., 2024; Einarson et al., 2024; Kazama et al., 2024).

One way to apply deep learning to animal movement is by representing the process as comprised of distinct but interacting behavioural processes, such as movement and habitat selection, analogous to a step selection function. Some benefits of using a deep learning approach include: 1) an ability to learn about spatial features of the landscape that may not be explicitly captured by the pixel values of the covariates, as we can use deep learning tools that were developed for image data; 2) covariate interactions can also be represented by flexible and non-linear processing layers, which can also interact with temporal dynamics over multiple time-scales; 3) given the modular capability of deep learning there is additional potential for extending the model by using the wide array of deep learning tools, such as recurrent layers (Drori, 2022) or transformers (Vaswani et al., 2017) to incorporate a history of previous locations, which may approximate memory processes; 4) the flexibility of deep learning also allows for other data sources such as accelerometers and physiological sensors to be integrated which could allow for a more holistic representation of animal movement behaviour (Williams et al., 2020; English et al., 2024).

In this paper we present an approach that formulates a deep learning model to replicate an SSF, with an explicit representation of movement and habitat selection processes. We term this approach deepSSF, which refers to the general approach, rather that any particular network architecture. There are numerous ways to formulate a deep learning model that could be used for modelling animal movement, and we hope to highlight one possible approach that may be readily extended to use other neural network architectures and incorporate more realistic processes such as memory and social dynamics. We show how the deepSSF approach may be used to learn about hitherto unknown behavioural processes underpinning the movement data, as well as for realistic predictions.

## 2 Materials and Methods

### 2.1 Study area and data collection

Data were collected from the Djelk Indigenous Protected Area in Western Arnhem Land, Northern Territory, Australia. The area is a culturally significant landscape comprised of tropical savanna with areas of open woodland, rainforest, a varied river and wetland system, and open floodplains. To understand their movement and habitat selection behaviours, 17 female water buffalo (*Bubalus bubalis*) were GPS-tracked in collaboration with Bawinanga Aboriginal Corporation rangers between July 2018 and November 2019. For further details on the data collection see Forrest, Pagendam, et al. (2024). To exemplify the deepSSF approach we arbitrarily selected a single individual’s data, which had 10,103 GPS locations that were suitable for training the model. There were 12 remaining individuals (*n* = 93, 446 GPS locations) that had high quality data which we used for out-of-sample validation.

### 2.2 Landscape covariates

Buffalos’ movement decisions are driven by factors such as vegetation composition and density for resource acquisition and shade, access to water, and the terrain (Campbell et al., 2020). In monsoonal ecosystems of Northern Australia, vegetation and the distribution of water changes dramatically throughout the year. To represent the seasonal changes in vegetation, we used monthly Normalised Difference Vegetation Index (NDVI), which measures photosynthetic activity and approximates the density and health of vegetation (Reed et al., 1994; Myneni et al., 1995). NDVI is an informative covariate in this landscape (Campbell et al., 2020) as it distinguishes between the broad vegetation classes, identifies wet and flooded areas, and can quantify buffalos’ forage resources as they are typically under open canopy. Monthly NDVI layers were generated from Sentinel-2 spectral imagery at 10 m x 10 m resolution using Google Earth Engine by taking the clearest pixels from a range of images for that month to alleviate the effects of obstruction from clouds. We selected the highest quality-band-score, which is based on cloud and shadow probability for each pixel, resulting in a single obstruction-free image of the NDVI values for each month of each year. We also used temporally-static layers for canopy cover and herbaceous vegetation, which are derivatives of Landsat-7 imagery and were sourced from Geoscience Australia at 25 m x 25 m resolution (Source: Geoscience Australia; Landsat-7 image courtesy of the U.S. Geological Survey). We represented broad-scale topographic features that affect buffalo movement by including a slope covariate, which was summarised from a 25 m x 25 m digital elevation model using the terra R package (Hijmans, 2024; R Core Team, 2024), and was calculated using the methodology of Horn (1981), which uses the elevation difference in the x and y directions of the eight neighbouring cells to estimate slope. The canopy cover layer was a proportion from 0 (completely open) to 1 (completely closed), and the herbaceous vegetation layer was binary, with 1 representing grasses and forbs, and 0 representing other (which is predominately woody growth). All spatial variables were discretised into grids (i.e. rasters) and resampled to be 25 m x 25 m resolution.

### 2.3 deepSSF model overview

Deep learning can be considered as a sequence of blocks, each of which perform some (typically nonlinear) transformation on input data to produce some output. Providing each block has the appropriate inputs, they can be combined to build a larger network that is capable of achieving complex and abstract transformations and can be used to represent complex processes. As such, we will describe the deepSSF model that we developed in terms of blocks that take some input, apply transformations, and produce an output. This description will also be helpful to understand the `PyTorch` code used for the model, which is itself comprised of blocks (called modules) (Paszke et al., 2019). Technical terms related to deep learning are summarised in the Glossary, found in the Supplementary Information, as well as a Supplementary section of deep learning concepts in the context of the deepSSF approach.

Our goal was to use a deep learning framework to generate next-step probability surfaces that can be sampled to generate animal movement trajectories. In our case we developed a model that had two distinct but interacting structures that we call subnetworks, which explicitly represented a movement process and a habitat selection process, respectively. Having processes that separately represent habitat selection and movement is analogous to a typical (parametric) step selection function (SSF), where there is a selection-free movement kernel that has distributions relating to how far the animal will travel in a given ‘step’ (step lengths) and how persistent its movement is (turning angles), as well as a habitat selection process that is described by a log-linear relationship to the surrounding environmental covariates (Avgar et al., 2016; Fieberg et al., 2021). In a similar way to how these two effects are combined in an SSF (Avgar et al., 2016), the outputs of the deepSSF subnetworks were then combined into the next-step probability. A similar approach of combining the outputs of neural networks was used by Pagendam et al. (2023), who termed it a ‘log-additive neural model’, as the outputs of the subnetworks are added on the log-scale.

In our deepSSF model, the primary model components were convolutional layers, which are key components in convolutional neural networks (CNNs) and are commonly applied to image data for classification, but can also be used for regression tasks (e.g. Pagendam et al. (2023)). Convolutional layers are often described as ‘feature-extractors’ (Hertel et al., 2015), as they determine salient features of images that affect the prediction outcome, and in our case the model could determine the features of the local landscape that influenced the animal’s movement. Convolutional layers are characterised by the use of convolution filters (also known as kernels) that sweep over the spatial inputs and apply element-wise operations, resulting in a set of ‘feature-maps’. The filters evaluate the local neighbourhood of grid cells, and with several successive layers they can represent features of the landscape such as forest edges, differing patch sizes, or riparian vegetation. The filters used by convolutional layers also have a depth component, meaning that interactions between all input covariates are considered, which can also include variables such as the time of the day and day of the year, as well as information about the animal such as its age or sex.

We describe each of the subnetworks in more detail below, and provide a conceptual overview of our deepSSF model in Figure 1.

**Figure 1:**
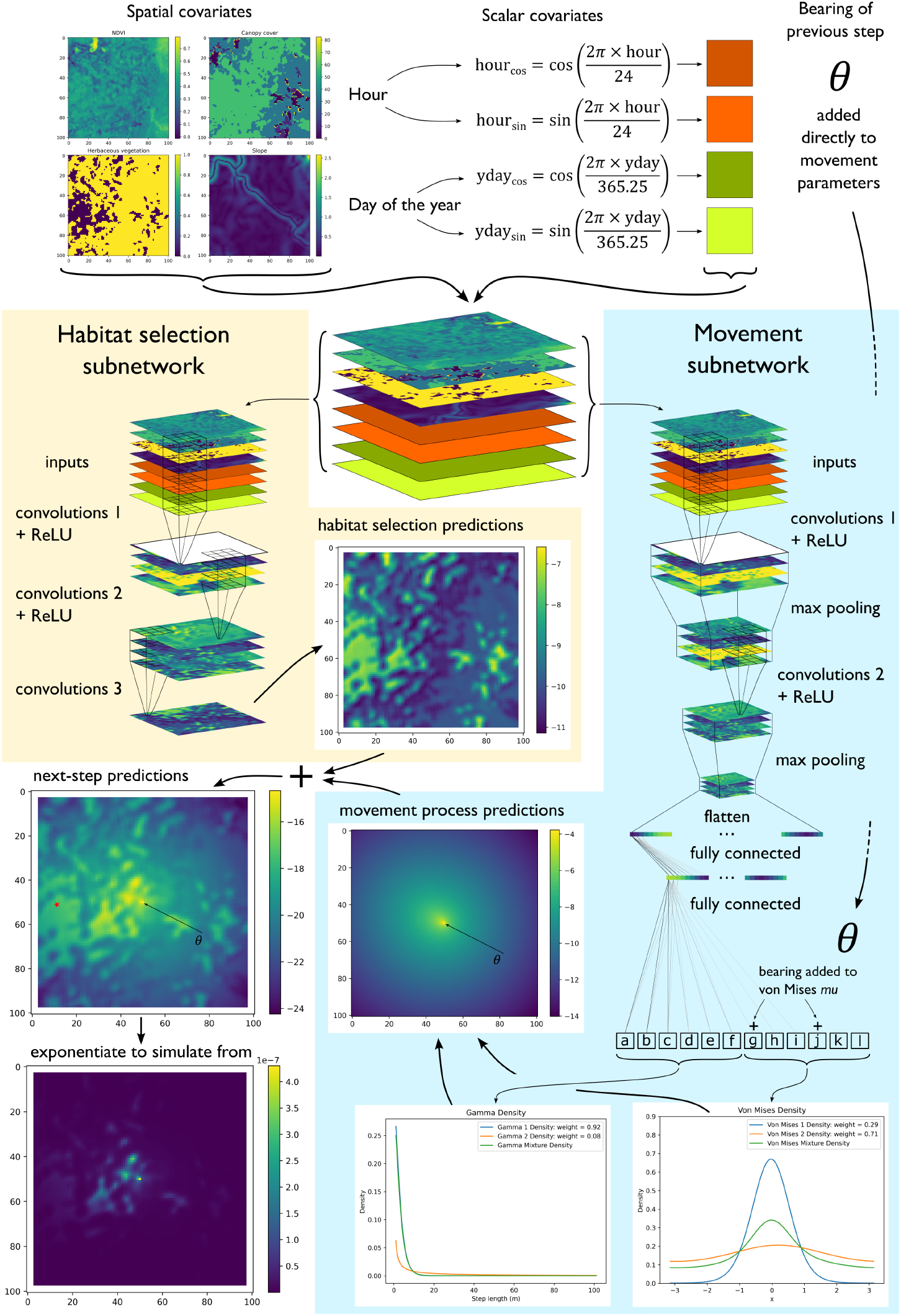
Conceptual overview of the deepSSF neural network used to predict animal movement. There are two subnetworks: a habitat selection and a movement process subnetwork. Both receive the same inputs, which are spatial layers such as environmental covariates, scalar covariates such as the hour, the day of the year (yday, also called ‘ordinal’ or ‘Julian’ day), and the bearing of the previous step. The periodic components (i.e. hour and yday) are decomposed into sine and cosine components to wrap continuously as a period, before being converted into spatial layers with constant values so they can be processed by the convolutional layers. To ensure that the turning angles are relative to the previous step, the bearing of the previous step is added directly to the predicted mean (*µ*) parameters of von Mises distributions. The habitat selection subnetwork uses convolutional layers that have parameters set to ensure that the output has the same spatial extent as the input, resulting in spatial, non-linear transformations of the input covariates, where all inputs can interact, to produce a probability surface (on the log-scale) describing the likelihood of moving to any cell based on the surrounding environment. The movement process subnetwork uses convolutional layers with max pooling to extract features from the input covariates that are salient to movement, and fully connected layers to process the convolutional layer outputs whilst reducing the dimensionality. The predicted output of the movement subnetwork can be any number of parameters that govern a movement distribution, so we used finite mixtures of two Gamma distributions for the step lengths and two von Mises distributions for the turning angles. This results in a total of 12 predicted parameters - a shape, scale and weight for each Gamma distribution and a *µ, κ* and weight for each von Mises distribution. The parameters are then converted to a two-dimensional movement probability surface (also on the log-scale) which is added to the habitat selection predictions, resulting in a next-step log-probability surface. To generate trajectories, the next-step log-probability surface is exponentiated and normalised such that the sum of all cells is equal to one, and a step is sampled according to these probability values. To highlight the directional persistence, the arrow and *θ* in the movement and next-step predictions denotes the bearing of the previous step, and the red star to the left of the next-step predictions is the location of the observed next step for those inputs.

### 2.4 Habitat selection subnetwork

The purpose of the habitat selection subnetwork is to transform the input covariates into a probability surface that describes the next-step probability whilst effectively ignoring the movement dynamics. To achieve this we used convolutional layers that apply transformations equally over the spatial extent of the inputs, producing ‘feature maps’ (for a conceptual overview of convolutional layers see the Supplementary Information). The habitat selection subnetwork therefore only had a single computational block which has three consecutive convolutional layers, the first two of which have rectified linear unit (ReLU) activation functions. The first two convolutional layers have four different 3 × 3 x *n* cell convolution filters that can each represent different features of the inputs, where *n* is the number of spatial inputs, resulting in four output layers (one for each filter). The final convolutional layer has a single 3 × 3 x *n* filter which acts to combine all input layers and therefore outputs a single layer, which is the habitat selection probability surface. The inputs for each convolutional layer were padded (see padding) with zeros and the filters had a stride of 1, which ensures that the spatial extent (i.e. number of cells) of the output of each convolution operation is the same as the input. This formulation allows for any number of convolution filters to be applied whilst retaining the same spatial extent. The number of convolutional layers and filters in our deepSSF model were chosen to be relatively low whilst still having enough complexity to represent the habitat selection process. Determining the number of convolutional layers and filters and the number of nodes in the feedforward network will vary by study due to the varying complexity of the data and the spatial layers. We suggest starting with a small model for faster training, and increasing it gradually until there are negligible improvements in the loss function on out-of-sample test data.

The convolutional layer can therefore be considered to apply spatial (as the filters consider neighbouring cells), non-linear (due to the ReLUs) transformations of the input data, where the probability values are trained to be higher where there are covariate features that were associated with observed next steps. Benefits of this formulation are that there is an interpretable intermediate output which is analogous to habitat selection of the input covariates, and importantly, it ensures that this subnetwork does not conflate with the movement process.

### 2.5 Movement process subnetwork

The purpose of the movement process subnetwork is to learn the intrinsic or ‘selection-free’ movement behaviour of the animal (Avgar et al., 2016), such that it can be used to inform the next-step predictions. As an animal’s movement behaviour is highly variable and stochastic, predicting it accurately for any particular step is an almost impossible task. However, an animal’s movement behaviour often follows patterns that can be described by common distributions, which may be mediated by factors such as the time of day. We therefore opted for a semi-parametric approach to describe the animal’s movement behaviour; instead of predicting the exact location of the animal’s next step, we predicted parameters governing step length and turning angle distributions that are likely to capture the animal’s movement dynamics for the next step. These parameters are then converted to a two-dimensional surface described by the predicted parameters, which becomes the output of the movement process subnetwork.

The movement process subnetwork that we used has three blocks, two of which are computational blocks with parameters that are trained, and one which converts the predicted movement parameters into a two-dimensional probability surface. Similar to the habitat selection subnetwork, the first block of the movement subnetwork was comprised of convolutional layers, except that for the movement process we also had max pooling layers after each of the convolutional and ReLU layers. We used a max pooling kernel size of 2 × 2 x n cells that had a stride of two, which means that the max pooling layer will reduce the total number of cells in the input grid by a factor of four, and retain only the most prominent features. We used two convolutional layers with four convolution filters each that were followed by ReLU activation functions and then max pooling layers. As there were two max pooling operations, the final feature maps had 16x fewer cells than the input layers. The size of the convolution filters were the same as in the habitat selection subnetwork (3 × 3 x *n*), although what the filters learn will be different, as in the movement subnetwork the filters will extract features from the covariates that influence an animal’s general movement tendencies.

To predict movement parameters, the dimension must be reduced so that the outputs are equal to the desired number of movement parameters. To achieved this we used a flatten operation on the outputs of the final layers from the convolutional block to a vector, resulting in a vector with a length equal to the total number of cells in the outputs of the final feature maps. We processed this vector using ‘fully connected layers’ (also called ‘dense’ layers), where every cell in the flattened vector connects to every cell in the next ‘hidden’ layer, which we gave a length of 128. These 128 cells are then connected to every cell in a final vector, which has a length equal to the number of movement parameters. To prevent overfitting, we used dropout at a rate of *p* = 0.1 (Srivastava et al., 2014) within the fully connected layers. As all cells in the fully connected layers connect, every cell from the initial covariate inputs can influence the final movement parameters, where the convolutional layers extract the most important features. The final block in this subnetwork takes the predicted movement parameters and converts them to a two-dimensional probability surface using the appropriate density function for each distribution.

As there can be any number of outputs from the fully connected layers, any form of step length and turning angle distributions can be used, whether parametric distributions or distributions described by other methods such as basis functions, providing that these parameters can be converted to a two-dimensional surface. This formulation then allows for finite mixtures of probability distributions, where there are multiple distributions that are combined together using a weighting. After an exploratory analysis of the movement characteristics, there appeared to be two modes in the step lengths (clearly visible after log-transforming the observed step lengths - Figure 4) and the turning angles were not described accurately with a von Mises distribution. We therefore fitted a mixture of two Gamma distributions for the step lengths and two von Mises distributions for the turning angles.

### 2.6 Next-step probability and loss function

To construct the next-step probability surface, the habitat selection and movement process probability surfaces are combined by adding them together when they are on the log-scale, in exactly the same way as for SSFs (Avgar et al., 2016). Prior being combined they are normalised whilst on the log-scale (using the log-sum-exp trick for numerical stability) to be valid probability surfaces (i.e. they sum to 1 after being exponentiated), so each subnetwork contributes equally to the next-step probability surface. We combined the normalised predictions of the habitat selection and movement processes prior to evaluating the loss function, which means that the final prediction accuracy depends on both subnetworks, and although they have separate sets of parameters and can learn their own representations, the subnetworks are trained simultaneously and implicitly depend on one another.

The loss function quantifies how accurately a model is predicting, which is used to update the weights and bias parameters of the network through backpropagation. We used a negative loglikelihood (NLL) loss function, where the target that the model is trying to predict is the observed location of the next step. For a 101 × 101 grid there are *N* = 10, 201 cells, each with a probability value, 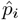 from the next-step probability surface. Each of these cells can be considered as a class with a predicted probability, and the value of the true class, 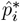, (i.e. the target, where the next step was observed to land) becomes the loss. The model is then trying to maximise this probability by minimising the negative log of this value. The negative log-likelihood for a single sample can be written as:

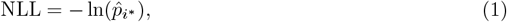

where 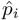 are the predicted probability values for all *N* cells where *i* = 1, …, *N* cells and *i*^*^ is the observed location of the next step. The next-step log-probabilities are given by

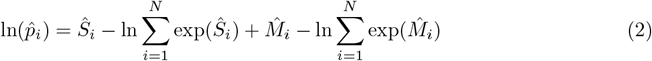

where *Ŝ*_*i*_ are the predicted values from the habitat selection subnetwork, which are outputted on the log-scale, and 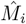 are the predicted values from the movement subnetwork, which are also outputted on the log-scale. Both of these surfaces are normalised with their respective normalisation constants, 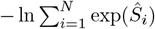 and 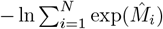 for the habitat selection and movement process predictions, respectively, such that they each sum to one when they are exponentiated. We use the log-sum-exp trick to calculate the normalisation constant, but we do not denote it here for clarity (a description can be found in the Glossary). Each set of inputs only has a single observed next-step, resulting in a single loss function evaluation.

### 2.7 Data preparation

To take advantage of deep learning tools that consider the spatial structure of the data, such as convolutional layers, the input data were formatted as grids (i.e. images or rasters). As we were generating next-step probability surfaces, we extracted the environmental information surrounding the animal’s current location, within the range of its typical step lengths. This resulted in a set of local rasters for each observed step. We chose a grid size of 101 × 101 cells, which is 2525 m x 2525 m, or 1262.5 m to the nearest edge of the local landscape. We chose this distance as 1262.5 metres comprises more than 95% of the observed step lengths, and it is important to have an odd number of cells so that there is a central cell, as that is where the animal is at the start of the step.

For the periodic inputs such as hour of the day and day of the year, to maintain their periodicity (so hour 24 is adjacent to hour 1, and yday 1 is adjacent to yday 365), we decomposed them into sine and cosine terms using *τ*_*sin*_ = sin(2*πτ/T*) and *τ*_*cos*_ = cos(2*kπτ/T*), where *τ* is the periodic covariate (the hour of the day or day of the year), and *T* is the largest value in the period, i.e. 24 or 365.25 (to account for leap years). To use convolutional layers which require images (i.e. grids or rasters) as inputs, we converted any scalar (single value) data to grids of the same spatial extent as the environmental covariates, and set the value of every cell to the value of the scalar.

To assess how well the model predicted the next step (i.e. to evaluate the loss function), we created an additional spatial layer corresponding to the ‘target’, which is the observed location of the next step. This spatial layer was comprised for all zeros, expect for a 1 where the next step was, so that when multiplied by the next-step probability surface, the sum of the resultant layer is the next-step probability where the observed step was.

### 2.8 Model training

The data were separated randomly into training, validation and test datasets, with an 80%, 10% and 10% split. As is typical in deep learning, the training data were split into batches (we used a batch size of 32 samples), and the loss function is evaluated for all samples in the batch. Using the average loss of a batch, the weights of the model are updated by performing backpropagation (Drori, 2022; Murphy, 2022), and then the next batch is put in. A complete iteration through all batches of the training data is termed an epoch. At the end of each epoch, the loss function is evaluated on all of the validation data, and if the average validation loss decreases the model is said to have improved (but the weights are not updated after evaluating the model on the validation data). The test data are reserved until training has completed. They are used to test the overall performance of the model on unseen data, which is typically used to compare between models.

The model was initialised with random weights, and we used the stochastic optimiser ‘Adaptive Moment Estimation’ (Adam) with an adaptive learning rate (the amount that the weights and bias parameters are updated) that decreases when the loss function plateaus. The learning rate started at 1*e* − 4, and decreased when the average validation loss had not decreased for five epochs, and training was terminated when the average validation loss did not decrease for 10 epochs. To prevent overfitting, the model weights are only saved when the validation loss decreased, so the final model weights are those that had the best performance on the validation data.

### 2.9 Simulating from the deepSSF model

Simulating from the deepSSF model requires selecting a starting location, hour of the day and day of the year (the bearing is usually randomly initialised). The local environmental covariates are cropped out as grids with 101 × 101 cells centred on the starting location, and the time inputs are decomposed into their respective sine and cosine components and converted to spatial layers. The trained model then processes the stack of spatial inputs, resulting in a next-step probability surface, which is exponentiated and sampled with respect to the probability weights. The sampled location is then the next step, and the process is repeated until the desired number of locations is reached.

We simulated 50 deepSSF trajectories that covered a three-month period of the late-dry season 2018, resulting in 2160 steps. We assessed whether the simulated trajectories displayed realistic characteristics of animal movement trajectories by visual comparison, and by comparing summary statistics of the movement and habitat selection with observed data. For the movement behaviour we compared the simulated and observed step lengths by plotting the full distribution of step lengths and turning angles, and also the mean step length for each hour of the day, which is a similar approach to Forrest, Pagendam, et al. (2024). For the habitat selection we binned the simulated and observed locations into hourly bins, and calculated the mean values of each of the covariates, which was also similar to (Forrest, Pagendam, et al., 2024).

### 2.10 Landscape-scale habitat selection

As trained convolution filters can be applied to images (i.e. rasters) of any size, it is possible to generate habitat selection surfaces of any size by using the trained filters from the habitat selection subnetwork. These have been trained to extract the salient features of the input data, and produce higher probability values where an animal is likely to take its next step, which also depends on the time of day and the day of the year. Additionally, these habitat selection filters have been conditioned on the movement process of the animal, as only local covariates that are ‘available’ to the animal are used in the training data, which was the reason for the initial development of step selection functions (Fortin et al., 2005; Rhodes et al., 2005).

When applied to a larger landscape area, the resulting probability surface denotes only habitat selection, and not the expected distribution of animals, as of course when this surface is generated it ignores the movement process, and assumes that an animal could access any point in the landscape at the next step. This is an assumption of resource selection functions (RSFs), and has been called a ‘naive’ estimation by Signer et al. (2017), because it ignores the movement dynamics. Predicting over broader areas should also be used with caution as covariate values that were not encountered during model training may produce inaccurate or misleading results. Acknowledging these assumptions, we include examples of landscape-scale habitat selection to suggest they have some utility. Primarily, the resultant habitat selection map provides a visual representation of what the habitat selection subnetwork has learned from the environmental covariates, and how this changes with respect to the other covariates such as the hour and day of the year. This can be used as a diagnostic tool for further model development, or as an end in itself, as it highlights the features present in the environmental covariates that the model considered to be influential in determining the next-step’s location. From these maps, we can also directly plot the correlation between the covariate values and the habitat selection prediction probability, which represents the marginal contribution of the covariate values (Supplementary Information Section Appendix C).

### 2.11 Next-step ahead validation and comparison to SSF

Whilst the realism and emergent properties of simulated trajectories are difficult to assess, we can validate the deepSSF models on their predictive performance at the next step, for each of the habitat selection, movement and next-step probability surfaces. Ensuring that the probability surfaces are normalised to sum to one, they can be compared to the predictions of typical step selection functions when the same probability surfaces are generated for the same local covariates. This approach not only allows for comparison between models, but can be informative as to when in the observed trajectory the model performs well or poorly, which can be analysed across the entire tracking period or for each hour of the day, and can lead to critical evaluation of the covariates that are used by the models and allow for model refinement.

We assessed the performance of the deepSSF model described above, as well as the performance of a deepSSF model trained with 12 Sentinel-2 bands at 25 m x 25 m cell resolution and slope as spatial inputs. We refer to this model as deepSSF S2, which is described in detail in the Supplementary Information (Section Appendix B). We compare these models to the performance of two typical SSFs fitted using conditional logistic regression to the same covariates as the deepSSF model. One of these models was fitted without temporal dynamics, denoted ‘SSF’, and the other was fitted with temporal dynamics on the movement and habitat selection processes over a daily period using two pairs of harmonics, as presented by Forrest, Pagendam, et al. (2024), which we refer to as ‘SSF 2p’.

We fitted each of these four models to a single buffalo’s data (ID 2005, *n* = 10, 108 steps), which was selected arbitrarily from individuals that had more than a year of high-quality data. We refer to this dataset as from the focal individual. We assessed the ‘in-sample’ performance of the models by generating and evaluating next-step-ahead predictions of the focal individual, and the ‘out-of-sample’ performance by using the focal individual model to generate next-step predictions for observed steps of the remaining 12 individuals that the model was not fitted to (*n* = 93, 446 steps).

### 2.12 Software packages

We used a variety of software packages to perform data exploration and processing, statistical analysis and model fitting, as well as model interpretation and plotting. In R (R Core Team, 2024) the primary packages that we used were `tidyverse` (Wickham et al., 2019) and `amt` (Signer et al., 2019) for data processing, `terra` (Hijmans, 2024) for processing raster data and `ggplot2` (Wickham, 2016) and `ggpubr` (Kassambara, 2023) for plotting. In Python the primary packages we used were `NumPy` (Harris et al., 2020) and `pandas` (McKinney et al., 2010) for data processing, `rasterio` (Gillies et al., 2013) for processing raster data, `PyTorch` (Paszke et al., 2019) for building and training the deepSSF models and `matplotlib` (Hunter, 2007) for plotting.

## 3 Results

On a Central Processing Unit (central processing unit - Dell Latitude 7420 with Intel Core i7 11th Gen 3.0GHz and 16 GB RAM) the deepSSF model took 1-2 hours to train, which typically comprised between 50-70 epochs (complete iterations through the training data), depending on the initial conditions. When trained on a Graphical Processing Unit (graphics processing unit) using Google Colaboratory (“Google Colaboratory”, 2024) the training time reduced to 10-15 minutes.

The deepSSF model was able to capture the habitat selection and movement dynamics of the buffalo’s data (Figure 2), resulting in simulated trajectories that represented important features of the data (Figure 3). The deepSSF approach generated trajectories that were visually similar to the observed data (Fieberg et al., 2024), although as the simulations did not have a memory or centralising tendency they wandered more widely than the observed data, and there was less obvious revisitation structure to the trajectories.

**Figure 2:**
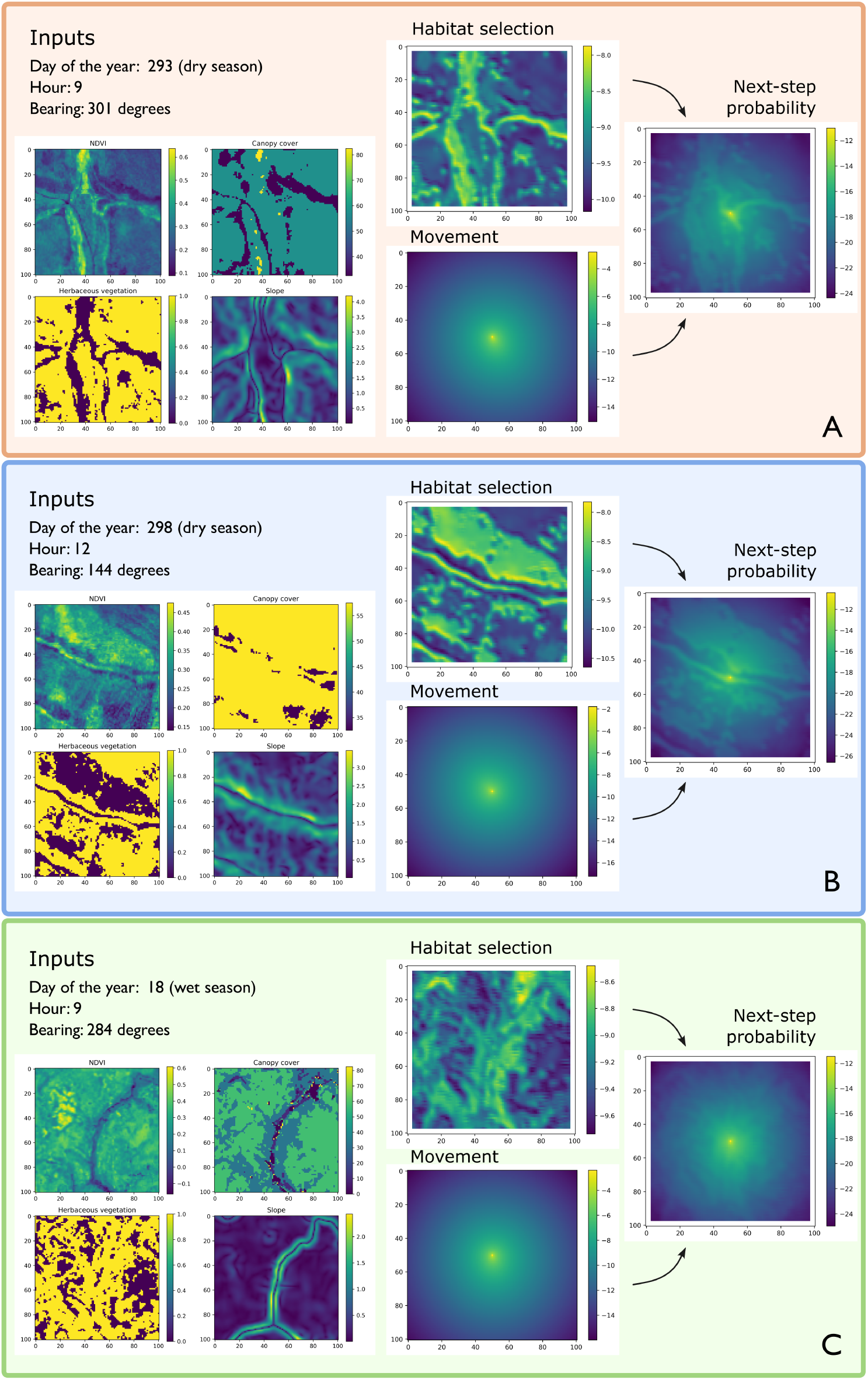
Here we show three example steps, with the associate inputs, intermediate predictions from the habitat selection and movement process subnetworks, and the resulting next-step probabilities outputted from the deepSSF model. The day of the year and hour inputs listed above the spatial covariates were decomposed into sine and cosine components and these values were converted to grids of the same spatial extent prior to inputting to the model. The bearing was added directly to the *µ* parameters of the von Mises distributions to deviate the turning angle from the previous step. The intermediate outputs provide an indication of what the model considers to be important for predicting the location of the next step. In Panel A, there is higher selection for the areas that have woody vegetation (when bottom left covariate is 0), with higher values of Normalised Difference Vegetation Index (NDVI), and low values of slope along a watercourse feature. In Panel B, there is high selection for the low slope feature running laterally (which is a watercourse), but also high selection for the southern edge of the woody area, possibly for refuge near the river. In Panel C, the covariate values are of similar magnitude, but the model identifies several areas in the north of the covariates as a high probability of selection, which are localised areas of floodplain that are know to be used by buffalo, particularly during the wet season.

**Figure 3:**
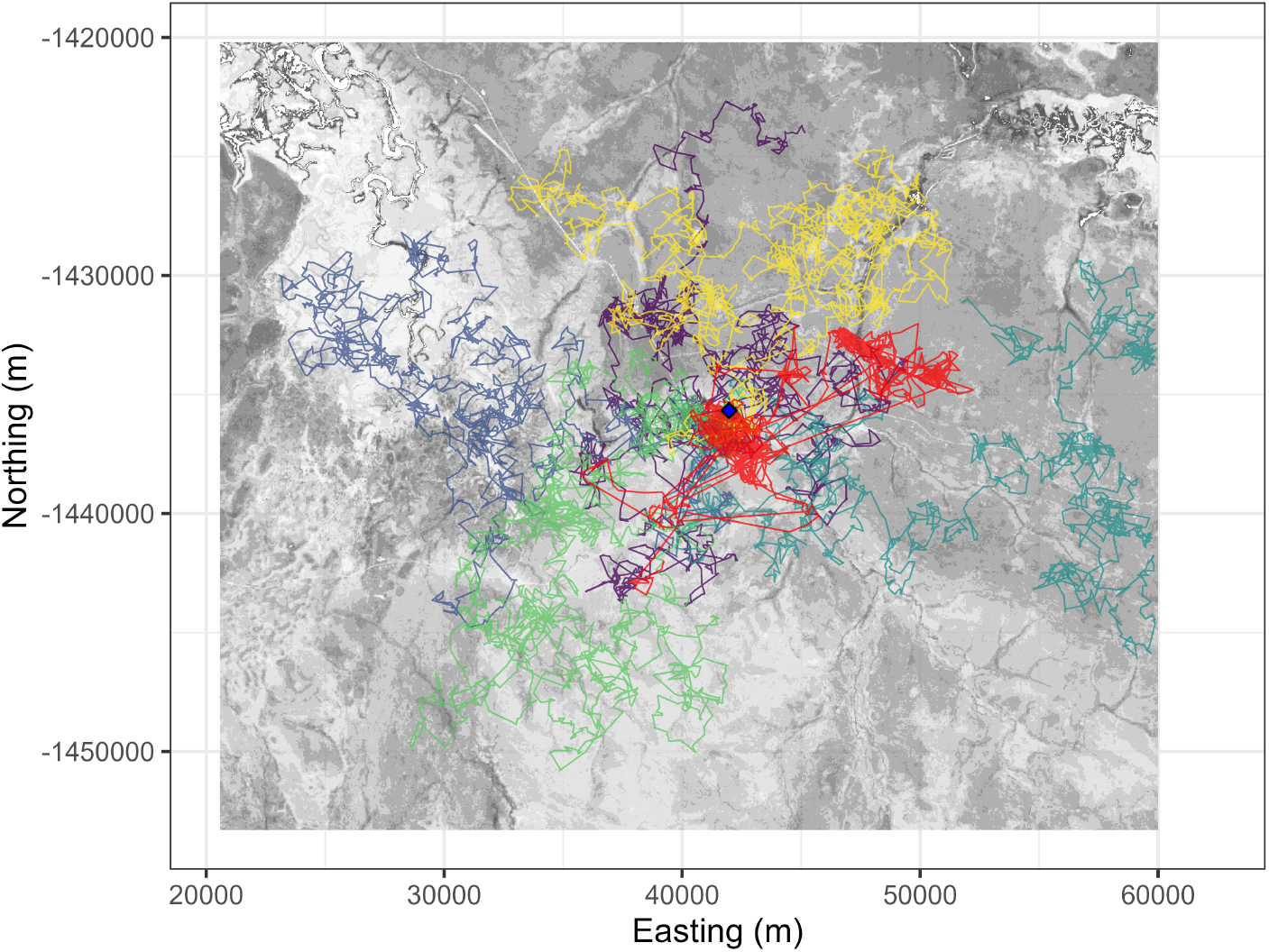
Example trajectories that were simulated from the deepSSF model, with Normalised Difference Vegetation Index (NDVI) as the background (with higher values as darker grey). The red trajectory is the observed buffalo trajectory, and the other colours are five simulated trajectories. Both the observed and simulated trajectories have 3,000 steps each. Without a memory process, the trajectories of the simulated data do not have any home ranging behaviour and wander more widely than the observed data.

The model clearly represented temporal dynamics in both the movement and habitat selection processes, which interacted over both hourly and seasonal time scales (Figures 4 and 5). When log-transformed, there was a clear bimodal pattern in the buffalo’s step lengths, which the deepSSF model was able to capture with the mixture of Gamma distributions. The observed turning angle distribution also did not follow a standard von Mises distribution due to the plateauing of turn angles away from the peak, but this was also captured well by the flexibility of the mixture of the von Mises distributions. The deepSSF captures the temporal dynamism in the observed movement dynamics, although does not replicate the peaks of movement around dawn and dusk as strongly (Figure 4). The deepSSF model was able to capture the hourly temporal dynamics in habitat selection, similar to the results of temporally dynamic step selection functions in Forrest, Pagendam, et al. (2024) (Figure 5).

**Figure 4:**
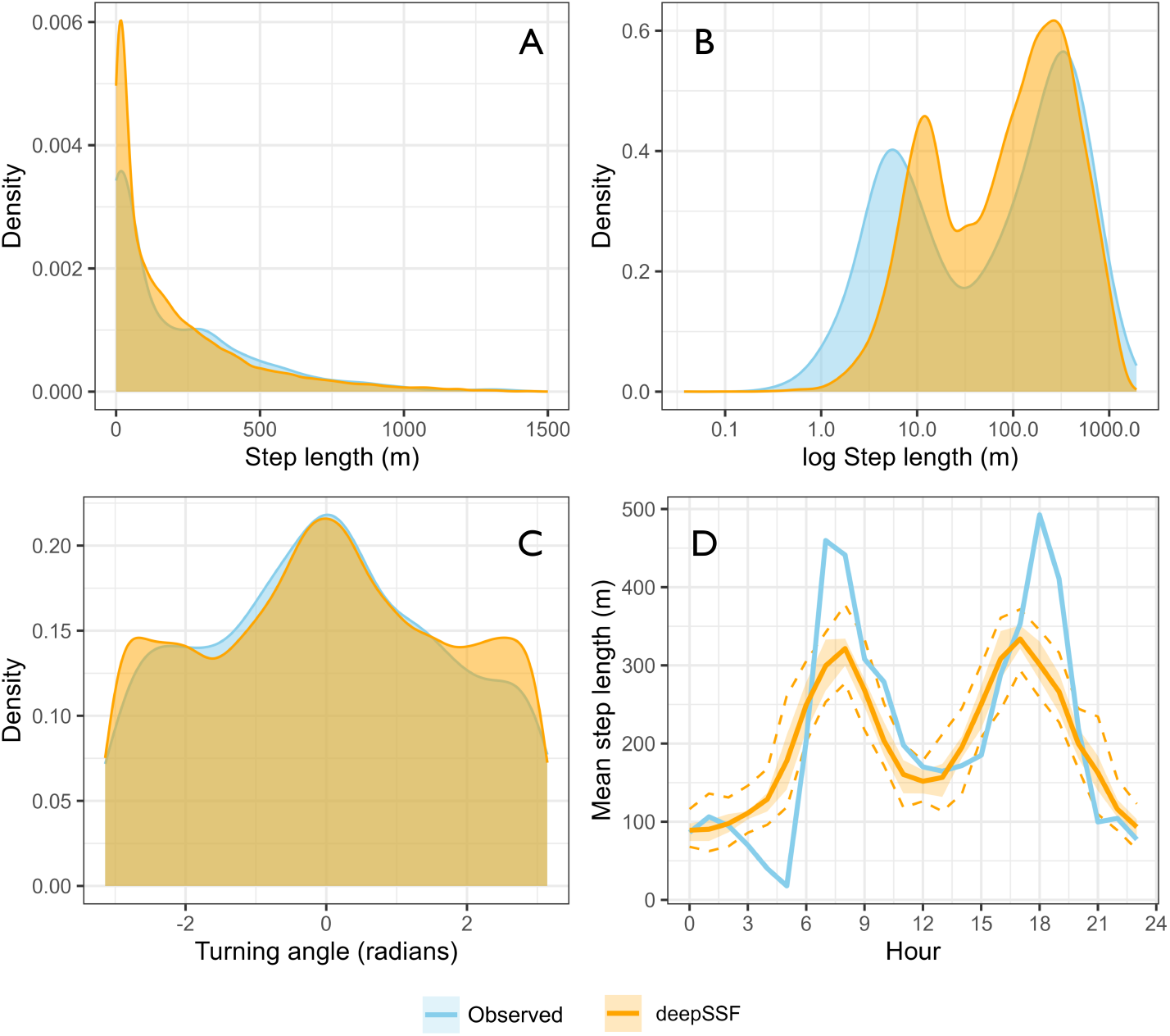
A comparison of the step length (panels A and B) and turning angle distributions (panel C), and the mean step length for each hour of the day (panel D), between the observed (light blue) and simulated trajectories (orange). We simulated 50 trajectories that covered a three-month period of the late-dry season 2018, resulting in 2160 steps. These simulated locations were compared against the observed movement data for that period. The mixture distributions predicted from the movement parameters in the deepSSF model were able to capture the bimodal pattern of the observed step lengths, and closely represented the observed distribution of turning angles, which is not able to be captured neatly by a standard von Mises distribution. The deepSSF captures the temporal dynamism in the observed movement dynamics, although does not replicate the peaks of movement around dawn and dusk as strongly as in the observed data.

**Figure 5:**
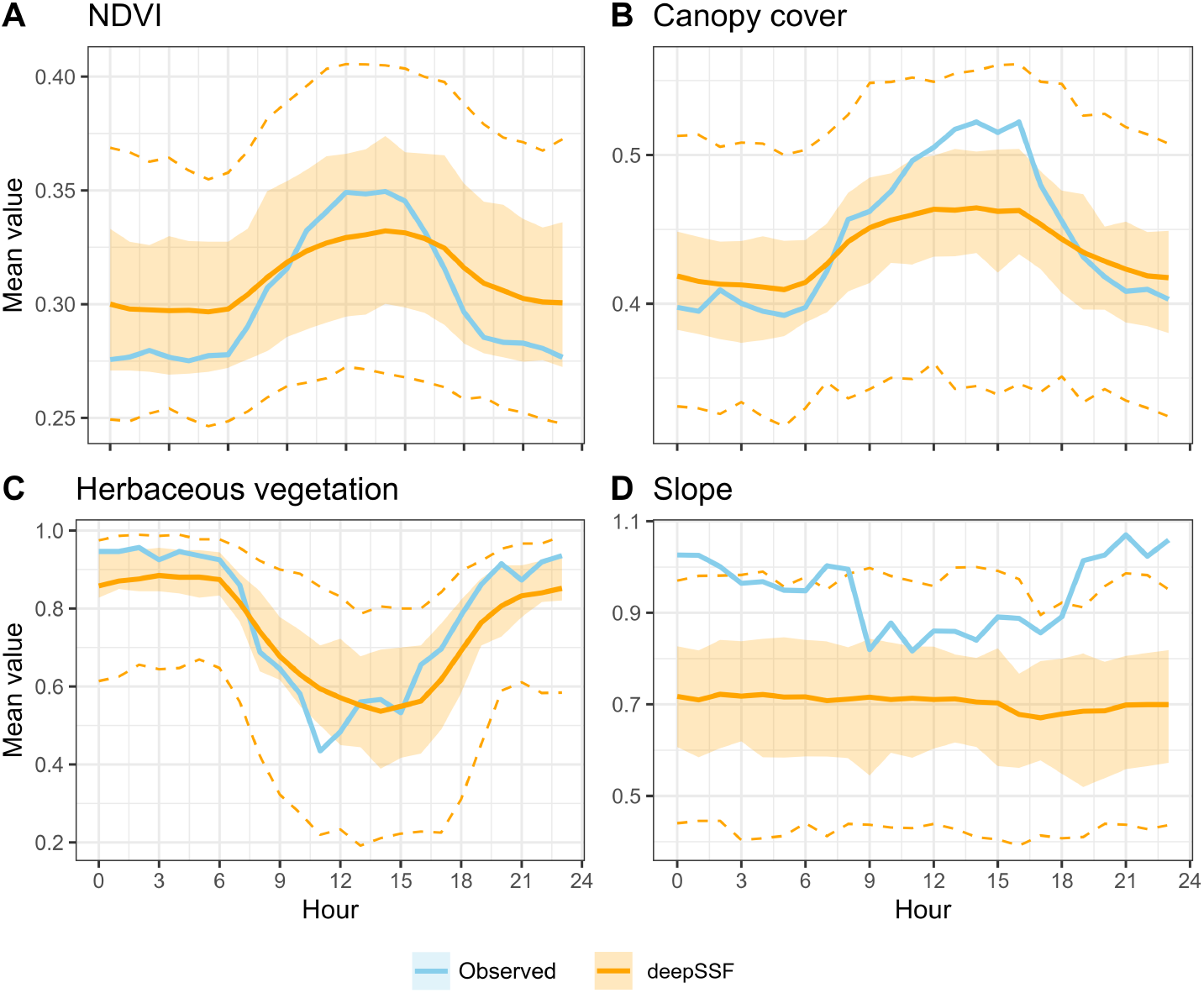
To assess the temporally dynamic habitat use of the simulated trajectories, we binned all simulated (orange) and observed (blue) steps into the hours of the day, and compared the mean values of each covariate at each hour of the day. We simulated 50 trajectories that covered a three-month period of the late-dry season 2018, resulting in 2160 steps, and we took a subset of the observed data to cover the same period. The deepSSF model was able to capture the hourly temporal dynamics in animal movement, similar to the results of temporally dynamic step selection functions in Forrest, Pagendam, et al. (2024). The shaded ribbons enclose the 25% and 75% quantiles, and the dashed lines are the 2.5% and 97.5% quantiles. The solid line is the mean for that hour across all trajectories or for the buffalo data. The deepSSF model captured the temporally dynamic habitat selection well for Normalised Difference Vegetation Index (NDVI), canopy cover and herbaceous vegetation, although the trend was not as clear for slope (although only a small range of slope values are shown on this plot).

The landscape-scale habitat selection maps clearly show the representation of hourly and seasonal dynamics of habitat selection (Figure 6). The habitat selection varies dramatically throughout each day, which for buffalo often has opposing trends at certain times of the day (Figure A7). Across the year the habitat selection also changes, which in this ecosystem is likely driven by the distribution of water (Figure 6).

**Figure 6:**
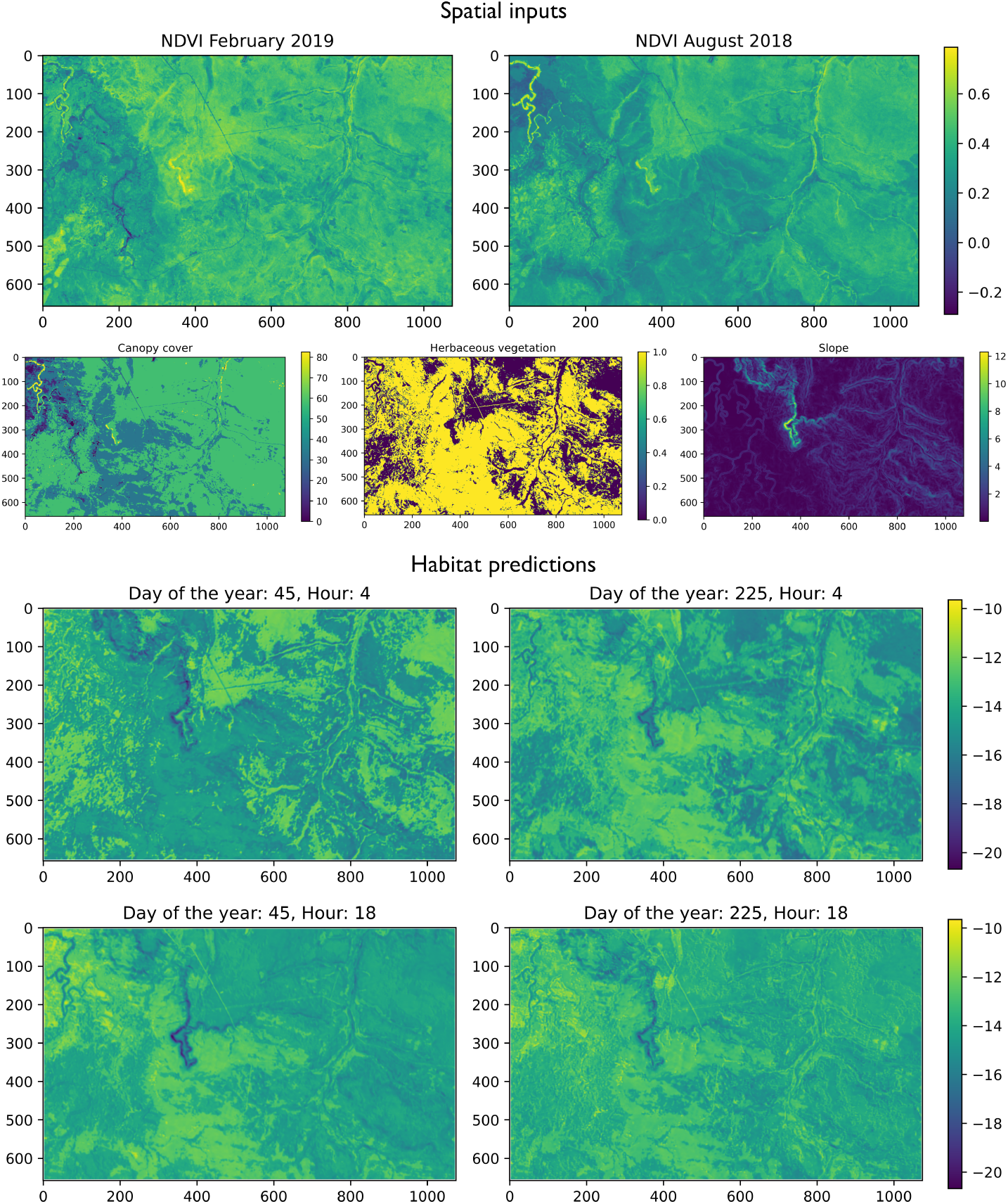
As convolution filters are fixed in size and apply transformations uniformly across grids, they can be applied to any landscape extent to approximate habitat selection (without accounting for movement dynamics). Here we show habitat selection at two times, 4am and 6pm, in the middle of the wet season (yday = 45) and the dry season (yday = 225). The spatial covariates are shown in the top panels, and the habitat predictions for each day and hour are shown below. These plots illustrate that the model was able to represent the multi-scale temporal dynamics of habitat selection that was present in the observed data. In the wet season, there was selection for higher canopy cover and woody vegetation in at Hour 4, and more open vegetation at Hour 18. In the dry season, there was more consistent selection for open habitat at these hours (but not during the middle of the day, which can be seen in Figure 2, Panels A and B).

When compared to typical SSF models, the deepSSF models had much higher average prediction values for the focal (in-sample) individual for the habitat selection, movement and next-step probabilities at the location of the next step, particularly for the deepSSF model fitted to 12 Sentinel-2 spectral bands and slope (Figures 7, A8 and A9 and Table 1). When trained on the derived landscape layers (i.e. not the Sentinel-2 data), the performance of deepSSF dropped for out-of-sample habitat selection predictions, showing weaker performance than either of the SSF models and was only marginally better than a null model, which is evidence of overfitting. However, when the deepSSF model was trained using Sentinel-2 data (deepSSF S2), the model’s in- and out-of-sample performance was much higher than both the deepSSF and SSF models. This suggests that these raw layers contain more information that is relevant to buffalo than oft-used derived measures like NDVI, particularly in the dry season. This was a similar result to when the probabilities were binned into the hours of the day, where the night-time performance of the deepSSF performance, but it was low during the day, again in contrast to the deepSSF S2 model.

**Table 1:** Next-step ahead probability values to compare between step selection functions and deepSSF models. We calculated the habitat selection, movement and next-step probability values for each step of the data that the models were fitted to (focal individual, a single buffalo - ID 2005), as well as 12 other individual buffalo that were not included in model fitting. The selection of the focal individual was arbitrary. Each of the prediction surfaces was normalised to sum to one such that they were valid probability surfaces, which makes the probabilities at the observed next step comparable between models. We compared two SSF models, a typical SSF that was fitted to the same covariates as above (NDVI, canopy cover, herbaceous vegetation and slope), as well as a temporally dynamic model with 2 pairs of harmonics across the hours of the day (SSF 2p). These models are the same as in Forrest, Pagendam, et al. (2024), except that they are only fitted to a single focal individual, and that they are fitted to all of that individual’s data, rather than being separated into wet and dry seasons. The deepSSF models are one that was fitted with the four spatial covariates listed above, as well as temporal dynamics for the hour of the day and the day of the year, and mixtures of two Gamma distributions for the step lengths and two von Mises distributions for the turning angles (deepSSF). The second deepSSF model was fitted directly to Sentinel-2 data, and also had temporal dynamics on the hour of the day and day of the year, and also had mixture distributions for the step lengths and turning angles (deepSSF Sentinel-2).

|  |  |  |  |
| --- | --- | --- | --- |
| *All values are $1 \times 10^{-4}$* | Focal individual ($n = 10,108$ steps) | | |
| Probability | Habitat | Movement | Next-step |
| *null* | *0.980* | *0.980* | *0.980* |
| SSF | $1.026 \pm 0.002$ | $17.08 \pm 0.141$ | $17.64 \pm 0.149$ |
| SSF 2p | $1.072 \pm 0.003$ | $27.27 \pm 0.333$ | $29.01 \pm 0.359$ |
| deepSSF | $1.841 \pm 0.025$ | $1262 \pm 25.16$ | $1289 \pm 25.67$ |
| deepSSF Sentinel-2 | **$2.296 \pm 0.025$** | **$1293 \pm 25.51$** | **$1339 \pm 26.41$** |
| *All values are $1 \times 10^{-4}$* | Out-of-sample (12 individuals, $n = 93,446$ steps) | | |
| Probability | Habitat | Movement | Next-step |
| *null* | *0.980* | *0.980* | *0.980* |
| SSF | $0.992 \pm 0.001$ | $17.38 \pm 0.047$ | $17.50 \pm 0.049$ |
| SSF 2p | $1.017 \pm 0.001$ | $26.58 \pm 0.105$ | $25.71 \pm 0.100$ |
| deepSSF | $0.984 \pm 0.003$ | $590.7 \pm 4.586$ | $523.7 \pm 4.308$ |
| deepSSF Sentinel-2 | **$1.391 \pm 0.006$** | **$772.6 \pm 5.830$** | **$718.5 \pm 5.702$** |

**Figure 7:**
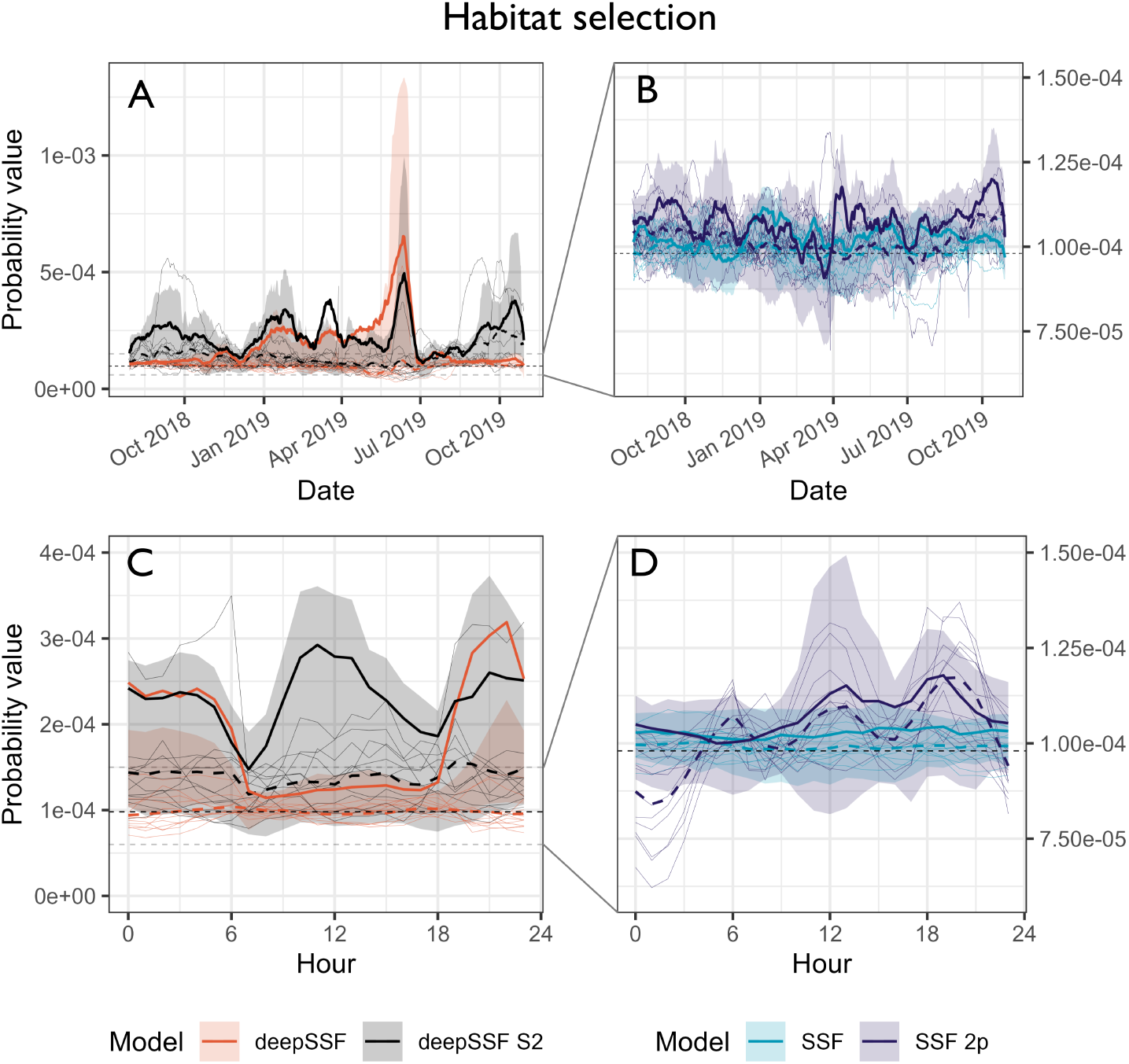
Probability values for the normalised habitat selection process at the observed location of the next-step. We compare between the deepSSF model with derived covariates (Normalised Difference Vegetation Index (NDVI), canopy cover, herbaceous vegetation and slope), the deepSSF model with slope and 12 Sentinel-2 bands at 25 m x 25 m cell resolution (deepSSF S2), a typical SSF fitted to the derived covariates, and an SSF with the derived covariates and temporal dynamics over a daily period using two pairs of harmonics, as presented by Forrest, Pagendam, et al. (2024). Panels **A** and **B** show the normalised habitat selection probabilities across the duration of the tracking period, smoothed using a moving window that was 15 days in duration and incremented by one day, and panels **C** and **D** show the normalised habitat selection probabilities across the hours of the day, where the probability values were binned into hour. The solid coloured lines show the average probability for the focal individual that the model was fitted to, and the shaded ribbon is the 50% quantile (there is high variability between probability values, so for clarity we omitted the 95% quantiles). The thin coloured lines are the average probability values for 12 individuals that the model was not fitted to, and are therefore out-of-sample validation data. The dashed coloured lines are the mean values for each model for all of the out-of-sample individuals. The SSF probability values were much lower than the deepSSF values (in both the positive and the negative direction), and the SSF plots in panels **B** and **D** are therefore zoomed in, as indicated by the dashed grey lines in panels **A** and **C**. For the focal individual at the broader temporal scale, the deepSSF and deepSSF S2 models both performed well between December 2018 and July 2019 (wet-season and early dry-season), although only the deepSSF S2 model performs well outside of this period, suggesting that the derived covariates lack information that is salient to buffalo during this period. This is also reflected when viewed at an hourly time-scale, where the deepSSF model performs well between 1800 and 0600 (during the night), although during the day only the deepSSF S2 model performs well, suggesting that the covariates do not describe what is important to buffalo in the middle of the day.

## 4 Discussion

In this paper we have presented an example of how deep learning can be applied to animal movement data, with promising results for generating realistic and accurate trajectories of animal movement, and potential for revealing subtle properties of animal movement behaviour. Deep learning is a highly flexible and data-driven approach, requiring minimal pre-emptive specification about the functional form of the model. This allows the model to learn its own representation about the nature of the movement and habitat selection processes represented by the deepSSF model, which lends a robustness to the model’s outputs. We have also shown that we do not necessarily require ‘big data’ for deep learning, here using only a single individual’s data with approximately 10,000 GPS locations (and of those the model was trained on only ∼ 8,000, with the remainder used for validation and testing).

As we expected, the deepSSF models outperformed the SSF models on the in-sample data, which was particularly the case for when the model was trained with Sentinel-2 spectral bands and slope as the spatial covariates (deepSSF S2). The performance dropped for out-of-sample data for all models (including SSFs), and the deepSSF trained with the derived covariates (NDVI, canopy cover, herbaceous vegetation and slope) performed worse than the SSF models, and was only marginally better than a null model, which bears some evidence of overfitting. However, the deepSSF S2 model, trained on ‘raw’ Sentinel-2 layers rather than derived quantities, retained greater accuracy than all other approaches for out-of-sample data, suggesting that these inputs contain more information that is relevant to buffalo movement and habitat selection than derived quantities like NDVI and slope. In general this suggests that, when using deep learning as a model for predicting animal movement, users should make as few *a priori* derivations as possible, instead providing the model with the rawest data possible and letting it find the relevant covarying derived quantities itself.

The value of using raw layers was particularly clear for predictions in the dry season and for the hours during the middle of the day, where the deepSSF model had a lower in-sample performance compared to the deepSSF S2 model. Assessing how the predictive performance of the models and covariates varies across different time periods is informative not only for the models’ overall performance which can be used for comparison and selection, but also for the relevance of the spatial covariates, which can be used to guide model refinement. However, the explainability of the model outputs when fitted directly to the Sentinel-2 data is lower, evidencing here a explainability-predictability trade-off (Dwivedi et al., 2023; Henriques et al., 2024).

The out-of-sample performance of the deepSSF model suggests that it would benefit from being fitted to more data to represent the broader environmental space that the buffalo population in this area is likely to encounter, rather than only seeing the restricted environmental space of a single buffalo. A model with a partial-pooling or hierarchical representation (i.e. a ‘mixed model’) may be particularly useful here, as the data from each individual would be considered as more dependent than the data between individuals (Muff et al., 2020). Until a solution is developed, a model can be trained on all of the observed data, without representation in the model that the data comes from different individuals (complete pooling), or separate models can be trained on the data from different individuals (such as we have here), which can each for creating a ‘population’ of individuals to simulate with. In addition to a limited environmental space, it is clear from the variable performance throughout time that there would be a similar trend when sampling outside of the temporal period of the data, highlighting the need for data collection across seasons, and at a fine enough resolution to capture variable patterns throughout the day. These results suggest that the potential for the deepSSF approach to be used for forecasting should be investigated, to explore how much data is required and under what environmental conditions (current and future) allow for generalisable predictions that can be extrapolated.

Despite providing useful information about the next step in a movement trajectory, the validation approach that we used did not assess the emergent features of the trajectories against the observed data. This consideration will depend on the underlying question of a study, but may relate to movement aspects such as a realistic home ranging or revisitation behaviour, accurate (dynamic) distributions in the landscape, or the generation of realistic connectivity pathways and movement corridors.

An additional limitation of the deepSSF configuration that we used is that it does not inherently represent parameter uncertainty, and only makes predictions conditional on the model parameters being equal to their maximum likelihood estimates. There are approaches to introduce uncertainty into the outputs (Harrison et al., 2024), and approaches that provide a more familiar parameter uncertainty quantification, such as Bayesian neural networks, that are undergoing promising development (Goan & Fookes, 2020), but are currently computationally demanding. If a model estimates parameter uncertainty, then trajectories can be generated from the distributions of estimated parameters, leading to higher variability in the predicted outputs. However, how much impact this will have on the overall predictions, and also how to interpret and present these results, is still undetermined.

Deep learning models are not interpretable in the same way as a simpler statistical model (i.e. we cannot understand the influence of particular parameters such as through coefficients), but in our case we have model explainability through visualising the intermediate (movement and habitat selection predictions) and final outputs (next-step predictions) from the model, summarising the trajectories, and from correlating the covariate values with the habitat selection probability values. From these model outputs we were able to determine that there is substantial daily variation, and some yearly variation, in buffalo movement and habitat selection behaviour, which corroborates with fine-scale dynamic SSFs (Forrest, Pagendam, et al., 2024) and with the observed data. The model also learned multi-scale temporal dynamics, with daily movement and habitat selection behaviour that also changed throughout the year. Whilst we acknowledge that understanding what the deepSSF model has learned requires some more investigation by the user, we also believe that because animal movement behaviour is a sophisticated process, the outputs of typical statistical models such as SSFs can oversimplify the process, potentially obscuring subtle insights into the behaviour of the species and the nature of the movement process.

There have been several other papers which have provided a conceptual foundation for this study. Dalziel et al. (2008) offered a similar approach to what we have presented here, although the study was not widely adopted, and since then deep learning has advanced rapidly. Cífka et al. (2023) used transformers to train a general model (MoveFormer) for animal movement, which also replicated a step selection function. This approach shows very promising results for representing memory by incorporating a history of locations and using a transformer architecture, although applying the model to a wide range of species demanded reducing the location frequency to two GPS fixes a day, ignoring fine-scale processes. This approach also required proposing random steps which are compared to the observed steps (as in SSFs), rather than directly inputting spatial information which can allow for spatial features to influence movement. Given the modular nature of deep learning, it is possible that components of the deepSSF approach and Moveformer may be combined to represent fine-scale behavioural processes while using the more flexible (but data-hungry) transformer architecture, and may reveal insights about the spatial memory process. Another promising approach for representing sequences is via (deep) reinforcement learning (Villeneuve et al., 2021), which may also be combined with other processing layers such as we have here (Chen et al., 2024), and can provide accurate movement predictions for several steps into the future.

We built the model to represent movement behaviour as distinct processes, each processed by separate subnetworks. Creating two subnetworks had several purposes and benefits - it created an analogue to an SSF, which is a widely used and trusted approach to modelling animal movement; it allowed for the intermediate outputs of the movement and habitat selection processes to be interpreted, improving upon the black-box nature of many standard neural network architectures; and it allowed for different processing layers, which enabled a semi-parametric approach when modelling the movement process by predicting parameters of common step length and turning angle distributions.

Whilst we present one possible deepSSF model here, there are a number of exciting possible developments that would lead to a more realistic representation of animal movement. Firstly, a straightforward extension is to retain the deepSSF architecture and processes that we presented, but to add more information to the inputs of the movement process (and/or into the habitat selection process), which may also include a representation of past locations (Schlägel & Lewis, 2014; Oliveira-Santos et al., 2016; Rheault et al., 2021; Ellison, Potts, Strickland, et al., 2024). In our configuration the model only received the bearing of the previous step (to confer turn angle persistence) as well as the surrounding spatial covariates. It would be straightforward to incorporate a greater number of previous turning angles and previous step lengths, which would confer a persistence in speed and approximate a momentum process, and may be particularly helpful in cases of highly correlated movement kernels, such as high-frequency tracking data or for animal migrations. An example of such a movement process is Wijeyakulasuriya et al. (2020), where 39 variables, such as lagged (*x, y*) coordinates and lagged *x, y* velocities, were used at each time step to predict the movement of ants.

Secondly, there are a number of tools in the deep learning toolbox that may be applied to extend the modelling approach we have presented here. Potential processing architectures may include recurrent layers or transformer layers which can accommodate long sequences as inputs, such as a history of locations, allowing the model to represent a flexible and data-driven memory process (Shenk et al., 2021; Cífka et al., 2023; Chen et al., 2024). An additional architecture that may be able to represent social dynamics may be graph neural networks (GNNs) (Zhou et al., 2020), where individuals in a population may be represented by nodes and their relationship to each other by edges, which can be parameterised from movement data (Scharf et al., 2016). A graphical approach may allow for predictive modelling at a population-level with a socially informed model across many individuals. Both of these processes (memory and social dynamics) may allow for more accurate predictions, but may also represent and reveal some of the more sophisticated and abstract mechanisms of these processes.

Thirdly, integrating other data sources, such as those from other biologging devices is likely to allow for a more nuanced and comprehensive representation of animal movement behaviour, that may also reduce the dependence on any particular temporal scale (DeRuiter et al., 2017; Leos-Barajas et al., 2017; Adam et al., 2019; Munden et al., 2021; Saldanha et al., 2023). This integration can be achieved due to the modular nature of deep learning, where a subnetwork may process sensor data such as from high-frequency accelerometers or magnetometers, which could be summarised to the same temporal scale as the locational data, and may for example become an input to the movement subnetwork, where the the level of observed activity would inform the movement kernel at the next step.

Finally, it may also be possible to train a ‘foundational’ or ‘general’ deepSSF model for animal movement from the data of many species, taking advantage of open-source databases such as Movebank (Kranstauber et al., 2011), similar to the approach of MoveFormer (Cífka et al., 2023). A model such as this could learn the principles of animal movement dynamics, which could then be used as a pre-trained model and ‘transferred’ to a species of interest (Pan & Yang, 2010). Besides representing general animal movement dynamics, a purpose of a model such as this would be that less data would be required for new species, enabling accurate predictions with few data.

## 5 Conclusion

Here we presented the deepSSF approach for modelling and predicting animal movement using deep learning. We built an approach that represented animal movement behaviour as distinct but interacting processes, each represented by a subnetwork. This approach allows for a greater modelling flexibility and modularity, and provides intermediate outputs that can be interpreted in direct reference to the input covariates. The deepSSF model we presented generated realistic trajectories, as well as provided insights into water buffalo’s daily and seasonal movement and habitat selection behaviour in Northern Australia. The deepSSF model fitted to derived covariates such as NDVI and canopy cover did not predict well to data outside of the training dataset, although the deepSSF model fitted to Sentinel-2 satellite imagery showed high predictive performance both in- and out-of-sample, across the full temporal range of the data. The results for both models will likely improve when more data is used for training, and be better able to generalise under different environmental scenarios.

We kept the processes in our deepSSF model straightforward to exemplify the approach, but there are numerous possibilities for extensions. There are a wide array of computational architectures that can represent different features of the data, processes such as memory and social dynamics may be represented, and almost any input data can be integrated by adding subnetworks, which may process different inputs and produce outputs but can be combined in latter stages of the model. Further developments are promising for high-frequency telemetry data and migration pathways, where a more complex movement process may better capture the highly autocorrelated structure of these data.

We consider the deepSSF approach to be a valuable addition to the toolbox for modelling animal movement, and we provide a combination of R and Python code to process data, train deepSSF models, and produce all outputs in this article (see online Supplementary Information). Due to the data-driven nature of deep learning, training a deepSSF model can represent and predict from the complex and abstract processes of animal decision-making on fine scales, and has the potential to reveal subtle insights into animal movement behaviour which may be difficult to ascertain from parametric statistical models.

## Code and Data Availability

Code to replicate all analyses, as well as information to help understand deep learning in the context of the deepSSF approach can be found at https://swforrest.github.io/deepSSF/, which is associated with the GitHub repository at https://github.com/swforrest/deepSSF.

## Author Contributions

SWF, DP, CH and AJH conceived and developed the deepSSF ideas; AJH designed the data collection; SWF and DP developed the modelling methodology; SWF analysed the data and led the writing of the manuscript. All authors contributed critically to discussions throughout the design and modelling process, to manuscript drafts, and gave final approval for publication.

## Acknowledgements

SWF was supported by an Australian Government Research Training Program Scholarship and a CSIRO top-up scholarship. CD was supported by an Australian Research Council Future Fellowship (FT210100260). This work was funded by the federal government’s Department of Agriculture, Fisheries and Forestry under the Control tools and technologies for established pest animals and weeds competitive grants program and the Smart Farming Partnerships Program (round 2). Computational resources and services used in this work were provided by the eResearch Office, Queensland University of Technology, Brisbane, Australia. SWF thanks Charlotte Patterson and members of the Applied Mathematical Ecology (AMEG) and Bayesian Research and Applications Group (BRAG) for helpful discussions and feedback on the work, and Maryam Goldchin of CSIRO for help setting up a Python environment.

## Generative AI statement

ChatGPT was used to generate a starting point for glossary entries, which were edited by the authors. GitHub Copilot was used during coding, predominately to generate straightforward and repetitive code. Generative AI was not used in the generation of the ideas or at any stage of writing of the manuscript.

## Conflicts of Interest

The authors declare no conflicts of interest.

## Glossary

### activation function

An activation function is typically applied to the outputs of each node within the hidden layers of a neural network to introduce non-linearity. Arguably the most common activation function is the ‘Rectified Linear Unit’ (ReLU), which sets any negative values to zero, and any positive values remain unchanged; ReLU(*x*) = max(0, *x*). Another common activation function is the logistic activation function, which maps values to a range between 0 and 1 using *σ*(*x*) = 1*/*(1 + exp^−*x*^).

### Adam

A widely used optimisation algorithm for training deep learning models, which are typically based on gradient descent and determine how the weights and biases of the neural network are updated given the gradients determined by backpropagation. Adam adjusts the learning rate for each parameter by estimating both an exponential moving average of the gradients (essentially a running estimate of the mean gradient) and an exponential moving average of the gradients’ squares, which approximates how much the gradients vary over time (an ‘uncentered variance’). This results in faster convergence and more stable updates than simpler gradient descent algorithms, particularly for problems with noisy gradients or sparse data, and is a common default optimiser in many deep learning frameworks.

### backpropagation

Backpropagation is the fine-tuning algorithm that calculates how each weight and bias in a neural network contributes to the prediction error described by the loss function (the difference between the network’s output and the true value). It then propagates this error backward through the network, informing how each parameter should be updated. Weights and biases that led to larger contributions of error will be updated more strongly.

### bias

The bias is an extra trainable parameter added to the weighted sum of inputs before the activation function is applied. It’s analogous to the intercept in a regression model, allowing the node to shift its activation boundary (for a ReLU, below which will be set to 0). Without a bias term, the node’s output would always pass through zero, restricting the network’s representational flexibility.

### block

A modular component of a neural network, in our case defined as a Python class (type of object with certain functionality described by its definition) inheriting from torch.nn.Module in PyTorch. A block encapsulates a sequence of operations, including layers (such as fully connected layers or convolutional layers) and activation functions, to process input data. Each block has a forward method (i.e. instructions) that defines the data flow through the network during inference or training.

### central processing unit

Serves as the primary processor in a computing system, orchestrating general-purpose tasks. In the context of deep learning, it is often responsible for managing data pre-processing, control logic, and task scheduling, although its performance is typically constrained by its sequential execution model, making it less efficient than GPUs for the high-dimensional matrix operations characteristic of neural network training.

### class

The target categories that a classification algorithm attempts to predict for given inputs based on training data.

### convolution filter

A convolution filter, also called a kernel, is a small, learned matrix (in our case 3 cells x 3 cells) that sweeps across a set of spatial inputs and applies element-wise operations to the input data. Through training, the filter adapts to capture specific features, such as edges or textures, from localised regions of the input. Each filter is optimised via backpropagation to extract important features from the spatial layers. For the deepSSF models the convolution filters give probability to habitat features that are associated with observed buffalo locations. See the Supplementary Materials for a more thorough description that includes illustrative figures.

### convolutional layer

A convolutional layer applies a series of learned convolution filters across the spatial dimensions of the input, producing feature maps that highlight important patterns or structures. This operation preserves spatial hierarchies within the data and allows the network to capture local dependencies. See the Supplementary Materials for a more thorough description that includes illustrative figures.

### dropout

To prevent overfitting, dropout is used to randomly drop a proportion of nodes and their connections from the model during training by setting them to zero, which happens on every forward pass of the model. As the model is essentially trained using many different subsets of the network, this ensures that certain pathways through the network do not become too specialised in representing some part of the process, and results in the learning being more equally distributed through the network. Dropout therefore helps generalisability to unseen data, as the model is not as reliant on certain overspecialised parts of the network.

### epoch

One complete pass of the entire training dataset through the neural network. The weights and biases are updated throughout each epoch, which should decrease the value of the loss function, and there are typically many epochs in model during.

### flatten

The flatten operation is often used to transform a three-dimensional tensor (output by a two-dimensional convolutional layer) so that it can be fed into a fully connected layer. The fully connected layer takes a vector (a one-dimensional tensor) as an input and so the elements of a three-dimensional, *d × d × F* tensor are simply rearranged into a vector of length *Fd*^2^ so that all numerical values are retained, but the dimensions of the object storing them has changed.

### fully connected layer

In deep learning a fully connected layer (also known as a dense layer) refers to a layer where each node (a computing unit with a weight, bias and activation function) is connected to every node in the preceding and succeeding layers. It is typically used in the final stages of a neural network to combine the features extracted by previous layers and produce the final output. See the Supplementary Materials for a more thorough description.

### gradient descent

A fundamental optimisation technique commonly used to train machine learning models (including neural networks). It works by iteratively updating the model’s parameters (weights and biases) in the direction that reduces the loss (or error). Specifically, gradient descent calculates the gradient (slope) of the loss function with respect to each parameter, then adjusts each parameter proportionally to that gradient, moving “downhill” toward a local or global minimum of the loss function.

### graphics processing unit

Specialized hardware units designed for parallel processing of large-scale data operations. In deep learning, they accelerate the training of neural networks by enabling concurrent execution of numerous matrix multiplications, which are central to both forward and backpropagation phases. This parallelism significantly reduces training time, particularly in convolutional neural networks (CNNs) and other large-scale models.

### log-sum-exp

When working with probability distributions we often want all values to sum to one, such that they are valid probability surfaces, and for the we need a normalising constant, which is the sum (or integral) of all the values. The log-sum-exp trick is a numerical technique that prevents overflow or underflow from very large or very small values. To calculate the normalising constant we have to exponentiate all values and sum them, which can produce very large values, resulting in overflow, or very small values, resulting in underflow, which are both due to the computer not being able to represent enough digits, which are then rounded. The normalising constant for log-probabilities is

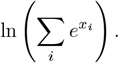

But what we can do is shift the values closer to reasonable numbers which will not cause over or underflow. For this we often use the largest value as a constant, *m* = max_*i*_ *x*_*i*_, and the expression becomes

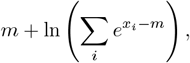

By subtracting the constant before exponentiating the values are kept within a safe numerical range, and the constant can be added back on afterwards to result in the normalising constant. This method is widely used in machine learning and statistics to stabilise calculations of log probabilities or normalisation constants. For a more in-depth description and examples see https://gregorygundersen.com/blog/2020/02/09/log-sum-exp/.

### loss function

Used to quantify the discrepancy between the predicted output of a neural network and the observed values (in this case the observed location of the next step). It provides a single value that reflects how well or poorly the model’s predictions align with the true observations. The objective during model training is to minimise this loss.

### max pooling

A dimension reduction operation commonly used in convolutional neural networks (CNNs) to reduce the spatial dimensions (height and width) of the input feature maps while retaining the most important information. It works by sliding a window (usually 2 × 2 or 3 × 3 cells) across the feature map and taking the maximum value from each window. By reducing the size of the feature maps, max pooling helps lower computational complexity, reduces overfitting, and provides some spatial invariance to small translations of the input. See the Supplementary Materials for a more thorough description.

### node

The fundamental computing units of neural networks, nodes have a weight, a bias, and an activation function that collectively transform input data into output signals. Each node’s operation is analogous to a simple regression, where the weight and bias are the model’s coefficients and intercept, respectively. However, unlike standard linear regression, nodes in neural networks typically also apply a non-linear activation function (e.g. ReLU or sigmoid). This added non-linearity helps neural networks represent complex patterns that can’t be captured with linear or logistic relationships.

### padding

Refers to the process of adding extra pixels (typically zeros) around the border of the input feature map before applying the convolution operation. The purpose of padding is to control the spatial dimensions of the output feature map. Without padding, the size of the feature map reduces with each convolutional layer due to the kernel’s movement across the input.

### recurrent layer

A type of neural network layer designed to process sequential data by maintaining a hidden state that is updated with each time step. Recurrent layers, such as Long Short-Term Memory (LSTM) and Gated Recurrent Unit (GRU) layers, are capable of capturing temporal dependencies in sequences by passing information from previous inputs to future outputs, making them particularly effective for tasks involving time series, natural language processing, and other sequential data.

### ReLU

A Rectified Linear Unit (ReLU) is a widely used activation function in deep learning, particularly in convolutional layers and fully connected layer. It introduces non-linearity into the model by transforming input values as follows: any positive input remains unchanged, while any negative input is set to zero, i.e., *f* (*x*) = max(0, *x*).

### stride

Defines how much the convolution filter (kernel) moves across the inputs. A stride of one means the filter moves one pixel at a time, while a larger stride means it moves by more than one pixel. Higher strides result in a reduced output feature map size, as fewer filter applications occur across the input. Stride impacts both the spatial resolution of the output feature maps and computational efficiency of the network, with larger stride being faster.

### target

Also known as the label or ground truth, it refers to the actual value or correct outcome that the model is trying to predict. It is the reference point used to measure the performance of the model by comparing the model’s predictions to the observed values through the loss function. In the case of the deepSSF model it is the location of the next step.

### tensor

A tensor is a general term for representing numerical data across multiple dimensions. A single number, known as a scalar, can be viewed as a rank-0 tensor. A list of numbers in one dimension—a vector—is a rank-1 tensor, and a rectangular grid of numbers—a matrix—is a rank-2 tensor. Beyond two dimensions, tensors simply extend this idea to higher ranks (e.g., rank-3, rank-4, and so on). This makes tensors highly flexible for representing complex data structures, such as images, time-series, and multi-channel signals, which often require more than two dimensions.

### transformer layer

A neural network layer that forms the basis of the transformer architecture, characterised by its use of self-attention mechanisms, which weigh the importance of different input elements, such as the preceding words in a sentence that give context to the current word, irrespective of their position in the sequence. Each transformer layer typically consists of self-attention and fully connected layers, enabling the model to capture complex dependencies in data, making it highly effective for tasks in natural language processing and time-series.

### weight

A weight is a trainable parameter that determines the importance of an input when generating an output. It’s analogous to a coefficient in a regression model: just as regression coefficients multiply the input variables, neural network weights multiply the input signals. In a fully connected layer, each input is multiplied by its corresponding weight before being summed (and offset by a bias); in a convolutional layer, weights define the convolution filter that scans over the input data. By adjusting these weights during training, the network learns patterns in the data that improve its predictions.

## Appendix A Deep learning concepts

### Appendix A.1 Convolutional Layers

The convolutional layers employed in our neural network architecture are also know as *two-dimensional convolutional layers*. Two-dimensional convolutional layers provide a mechanism to learn new features or variables of predictive value from gridded data using sets of convolutional weights that are re-used at different locations over two-dimensional space. The input to a two-dimensional convolutional layer is a three-dimensional input tensor, ***Q***, with dimensions (*d, d, n*). In the context of our application, ***Q*** represents a stack of *n* spatial covariates in a *d × d* pixel region of habitat surrounding an individual, which in our case was 101 × 101 cells with 25 m x 25 m resolution. Each of the *n* spatial covariates (i.e. NDVI, slope, etc.), is usually referred to as a ‘channel’. The convolutional layer transforms this stack of spatial layers into a new three-dimensional tensor, ***O***, with dimensions (*d, d, F*) through the application of *F* unique convolution filters (also called kernels) that are learned in neural network training. As convolution layers can be considered to transform the inputs covariates and ‘extract features’ from them, the resulting layers denoted by *F* are often called ‘feature maps’ (Figures A1 and A2). Each filter can be represented as a (2*w* + 1) *×* (2*w* + 1) *× n* tensor of weights and we denote the collection of these filters as ***W*** ^(1)^, …, ***W*** ^(*F*)^. The hyper-parameters *w* (the spatial width and height in pixels of the filter) and *F* are user-specified, and in our application were set as *w* = 1 and *F* = 4 in all cases except in the final convolutional layer that had *F* = 1 to aggregate the preceding feature maps into the final habitat selection map (Figure A2). For *w* = 1, each filter can be thought of as consisting of *n* convolution filters of dimension 3 *×* 3. Applying filter *f* ∈ {1, …, *F*}, results in a *d × d* matrix, ***O***^(*f*)^, with elements

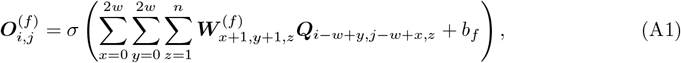

where we take ***Q***_*i,j,z*_ = 0 for any index for which one or more of the following is true: *i <* 1, *i > d*_*i*_, *j <* 1, or *j > d*_*j*_. The parameter *b*_*f*_, denotes the added bias parameter (a scalar that is analogous to an intercept in a regression model) for each filter and that is also learned during training. The function *σ*(·) is an activation function that is applied element-wise to its argument. For our model, the activation function was the rectified linear unit (ReLU), defined as ReLU(*x*) = max(0, *x*). The output tensor of the two-dimensional convolutional layer is the tensor created by combining each of the matrices ***O***^(1)^, …, ***O***^(*F*)^ along a third dimension. The elements of ***O*** can therefore be written as 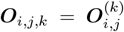. Ultimately, each two-dimensional convolutional layer introduces *F* [(2*w* + 1)(2*w* + 1)*n* + 1] parameters to the model that must be learned from the training data.

**Figure A1:**
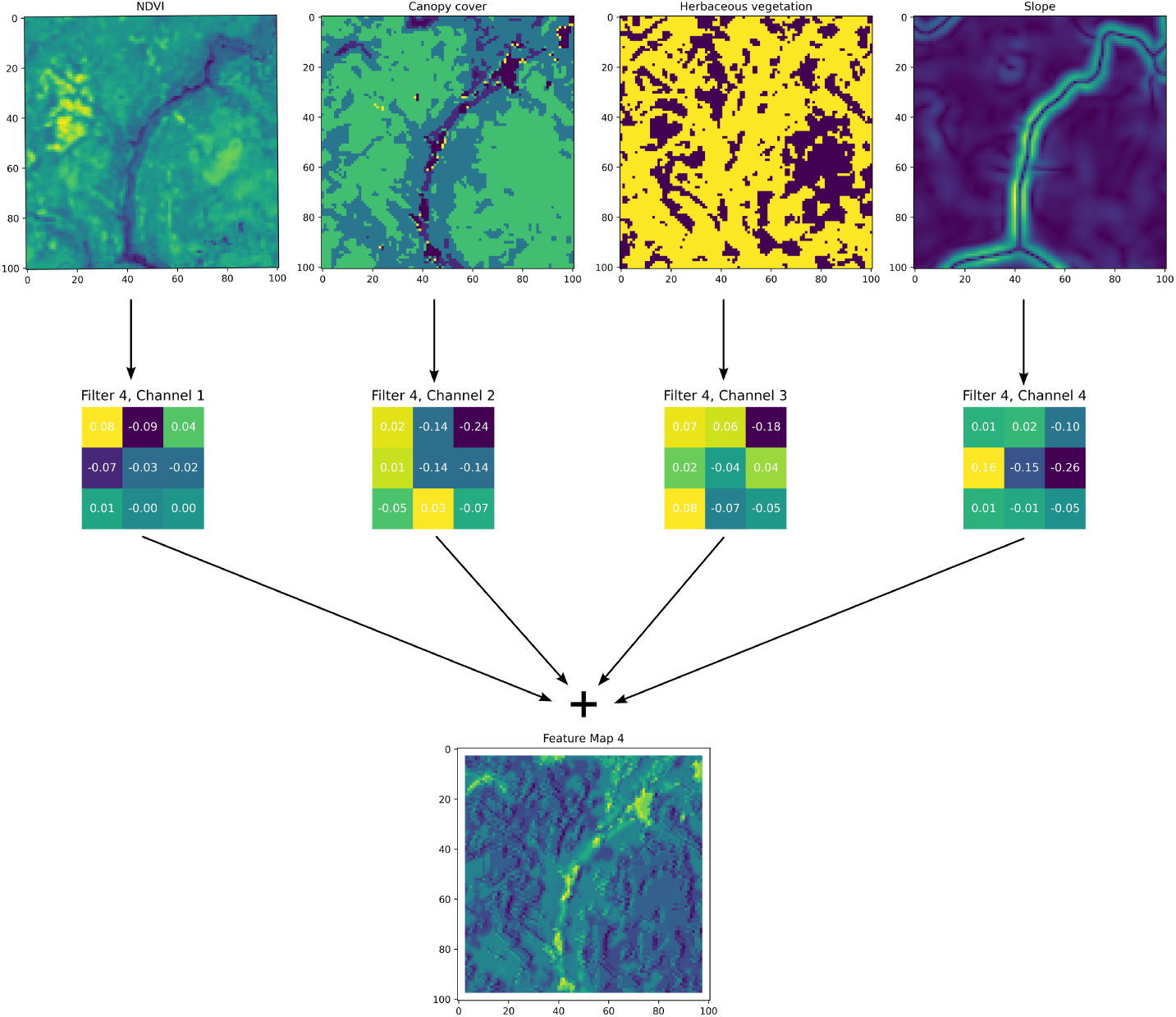
Here we show a **single** convolution filter from the habitat selection network and the resulting feature map (prior to being processed by an activation function, which in our case was the ReLU). A single convolution filter has *n* channels that relate to the *n* channels of the input layers, and each filter channel convolves over its respective input layer, which are then added together to create the feature map for that filter. To understand the convolution process, imagine the convolution filter channel starting in the top left corner of the spatial input, say NDVI, with the central cell of the filter in the top left corner. For each cell, the value of NDVI will be multiplied by the corresponding cell in the filter channel, resulting in nine values, which are then summed together to form the top left cell in the *intermediate* feature map (which are not shown here). The filter then moves to the right by one cell (as the stride is equal to one), and the element-wise multiplication is repeated. This results in transformations of many small, overlapping windows across the spatial input, resulting in another layer (the intermediate feature map). This process is repeated for the next input layer, which has its own filter channel, until each input layer has been processed into an intermediate feature map. These intermediate feature maps are then summed to create the output feature map for that filter. During training it is the values of the convolution filters that are updated. It can be helpful to see this process animated, and an example of an animation of a convolution filter across multiple channels can be found at: https://animatedai.github.io/. In the linked animation, the inputs are 7×6 ×8, and each coloured block in the middle represents a 3× 3× 8 filter, and in total there are 8 filters. Each coloured layer in the output to the right represents a feature map. Note that there is no padding for these operations, which results in a dimension reduction, but below there is an animation is shown with padding which retains the *d × d* dimensions of the inputs, such that we have used in our deepSSF model.

**Figure A2:**
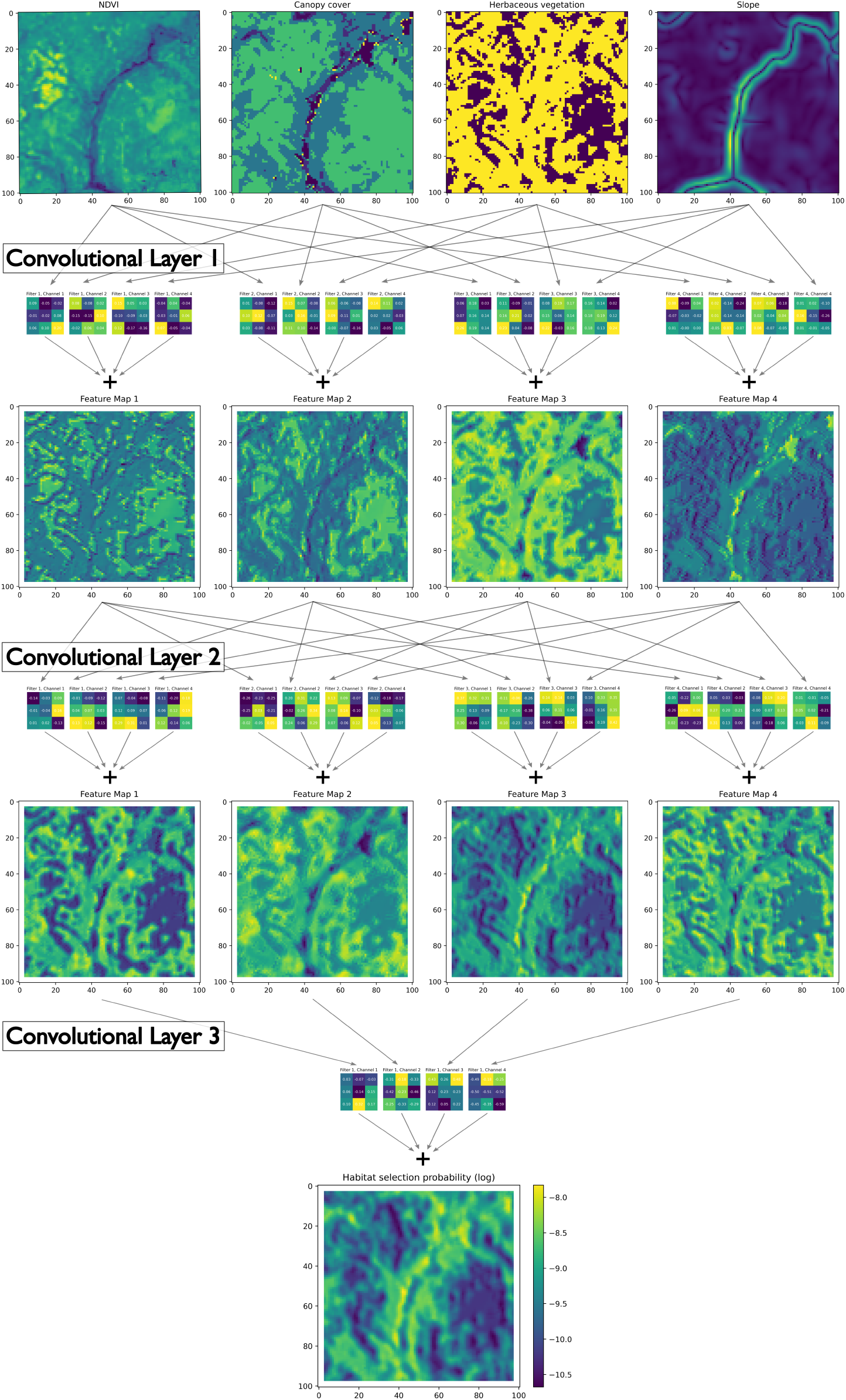
The benefits of deep learning come from many small operations combining together to represent complex and abstract processes. Here we show all of the convolutional layers that were used in the habitat selection subnetwork, except that for clarity we do not show the temporal inputs (hour of the day and day of the year decomposed into sine and cosine terms), which are also converted to spatial inputs, as shown in Figure (1) in the main text. Each convolution filter has a channel for each of the spatial inputs and produces a single feature map. In Convolution Layer 1 there are four filters, resulting in four feature maps, which each extract different aspects and features of the inputs (i.e. transform them in different ways). In Convolutional Layer 2 there are also four filters, except that the inputs to this layer are the feature maps outputted by Convolutional Layer 1. Convolutional Layer 3 then takes the feature maps from Convolutional Layer 2 as inputs and processes and aggregates then into a single feature map, which are the log probabilities of the habitat selection subnetwork (there is no ReLU after this layer meaning that there can be values less than zero). The successive convolutional layers are what give deep learning their name, where ‘depth’ is described by the number of successive processing layers, allowing for the models to learn a representation for abstract features in the input covariates.

### Appendix A.2 Max Pooling

Max pooling is an operation that is often used in convolutional neural networks to reduce dimensionality and condense information. Here we present a typical way that max pooling is applied in practice (and how it is applied in our model). The max pooling operation is best thought of as a function that accepts an *a × a* matrix, ***A***, as input and outputs an *a*^⋆^ *× a*^⋆^ matrix ***A***^⋆^, where 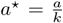. For simplicity here, we assume that 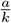 is integer-valued, but where this is not the case, the dimensions of the matrix ***A*** can be modified using “padding” to augment the matrix with additional rows and columns at it’s edges.

We refer to *k* as the kernel-width in the max pooling operation and use this to partition the matrix ***A*** as

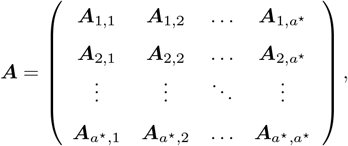

where each sub-matrix ***A***_*i,j*_ (*i, j* ∈ {1, …, *a*^⋆^}) is of dimension *k × k*. The output of the max pooling operation is the matrix ***A***^⋆^ which we define as

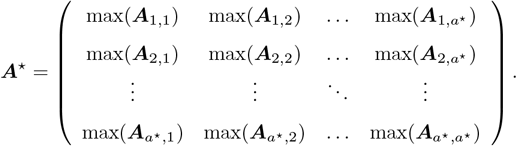

Max pooling can therefore be seen as simultaneously reducing the dimension of a matrix by a factor of *k*, and highlighting the extremes of the input feature matrix ***A*** over space.

In practice, max pooling operations often follow two-dimensional convolutional blocks in a neural network model. Where the convolutional blocks have more than one filter, the input to the max pooling operation will actually be an *a × a × F* tensor (a three-dimensional array), where *F* is the number of filters that are learned by the model in each convolutional layer. In this case, the max pooling operation described above is simply applied independently to each of *F* matrices of size *a × a* and each of the *a*^⋆^ *× a*^⋆^ sub-matrices are stacked to create a *a*^⋆^ *× a*^⋆^ *× f* tensor that is passed to the next layer in the network (often another convolutional layer).

### Appendix A.3 Fully Connected Layers

A fully connected layer in the neural network, maps an input vector ***u*** of length *n*_*u*_ to an output vector ***o*** of length *n*_*o*_ using

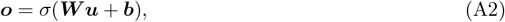

where *σ*(·) is referred to as the activation function and is applied element-wise to the argument. The matrix ***W*** is an *n*_*o*_ *× n*_*u*_ dense matrix of weights (with elements ***W***_*i,j*_ ∈ ℝ) that connects every element of the input vector to every element of the output vector, and ***b*** is a vector of length *n*_*o*_ (with elements *b*_*i*_ ∈ ℝ) and is called the bias vector of the layer. Each fully connected layer in the neural network introduces *n*_*o*_(*n*_*u*_ + 1) parameters that must be learned from the training data.

In our neural network architecture, the activation function is usually taken to be the rectified linear unit (ReLU) which is defined as ReLU(*x*) = max(*x*, 0). An exception to the use of ReLU in our model is where a fully-connected layer outputs a value that is taken to be the parameter of a density function. Our model has fully connected layers that predict the parameters of: (i) a two-component mixture of Von Mises densities; and (ii) a two-component mixture of Gamma densities. The former is used to model the direction for the next step in the animal’s trajectory and has parameters *p*_1_, *p*_2_, *κ*_1_, *κ*_2_, *µ*_1_, and *µ*_2_. The latter is used to model the step length of the next step in the trajectory and parameters relating to the mixture density, *p*_3_, *p*_4_, as well as shape parameters denoted *α*_1_ and *α*_2_ and scale parameters denoted *β*_1_ and *β*_2_ respectively. Each of these parameters have important bounds and we therefore use a different activation function to ensure that these bounds are honoured. For the parameters that can take any value in (−∞, ∞) (i.e. *µ*_1_ and *µ*_2_), the activation function used is simply the identity function: *σ*(*x*) = *x*. For the parameters that are required to be greater than zero (i.e. *α, β, κ*_1_, and *κ*_2_), the activation function used is the exponential function: *σ*(*x*) = exp(*x*). For the mixture proportions, *p*_1_ + *p*_2_ and *p*_3_ + *p*_4_ must each sum to exactly 1, so we normalised them via *p*_1_ = *p*_1_*/*(*p*_1_ + *p*_2_), *p*_2_ = *p*_2_*/*(*p*_1_ + *p*_2_) and *p*_3_ = *p*_3_*/*(*p*_3_ + *p*_4_), *p*_4_ = *p*_4_*/*(*p*_3_ + *p*_4_). As one *p* can be identified from the other, we could have also used a single *p* for each mixture density, and normalised it using the logistic activation function *σ*(*x*) = 1*/*(1 + exp(−*x*)) to be in the interval [0, 1], and then used 1 − *p* for the remaining mixture proportion.

## Appendix B Training a deepSSF model with Sentinel-2 imagery

Another data type that deep learning can accommodate is satellite imagery (e.g. red-green-blue (RGB) other spectral bands). Satellite imagery is a rich source of information that is clearly interpretable to us as humans, providing information about a range of environmental factors. Spatial layers such as Normalised Difference Vegetation Index (NDVI) and Normalised Difference Water Index (NDWI) are derived, often with simple arithmetic operations, from remotely-sensed images such as from the spectral bands of the Sentinel-2 satellites. Whilst using layers such as NDVI have useful interpretability benefits when fitting regression models, e.g. a higher habitat selection coefficient for NDVI suggests that an animal selects for higher environmental greenness, allowing a model that is flexible enough to learn its own representation and transformations of the raw spectral bands may lead to improved predictive performance.

However, the benefits of fitting a deepSSF model to satellite imagery directly may not only be for predictive performance. As we typically interpret satellite images using a true colour representation from the red-green-blue (RGB) spectral bands, the can examine the model’s predictions against the RGB bands, as that is part of the information the model is using to generate predictions. A researcher may easily identify features of RGB layers, such as certain vegetation types that may be not be interpretable from layers such as NDVI, and the predictions of the model can be assessed against these features. Additionally, this allows for the fitting of deepSSF models to the imagery from other remote sensing approaches such as aerial imagery or drones, which typically only capture true colour (RGB) images.

To assess the performance of the deepSSF approach using raw satellite images, we fitted the deepSSF model using 12 Sentinel-2 spectral bands, as well as the slope layer described in Section 2.2, as the spectral bands do not directly capture information about the terrain. The model structure was the same, except that there were 13 spatial layers (12 spectral bands and slope) and the same four layers representing time (a sine and a cosine term for the hour of the day and a sine and a cosine term for the day of the year), and the relevant inputs and outputs of the different neural network layers were updated such that the model could run.

## Appendix C Additional deepSSF model outputs

**Figure A3:**
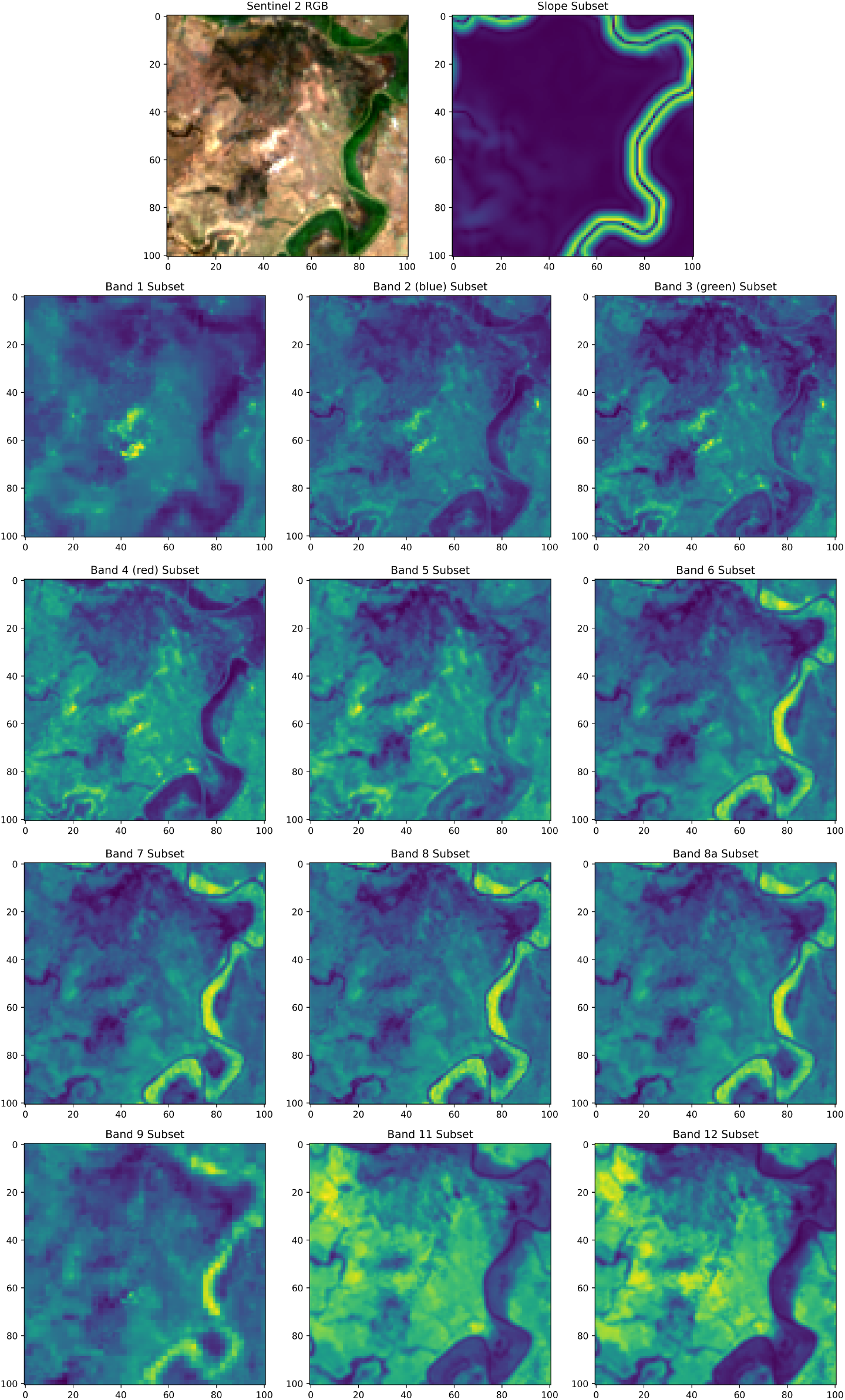
Example of spatial inputs (101 × 101 cells) for the deepSSF model trained on the 12 Sentinel-2 bands (deepSSF S2) for a single step. A true colour visualisation of the images is shown in the top left, which is from the Red (Band 4), Green (Band 3) and Blue (Band 2) bands, often termed simply ‘RGB’. The slope covariate, which was used in training the deepSSF S2 model, is shown in the top right. Four of the bands that we used have a native resolution of 10 m x 10 m cells (B2, B3, B4 and B8), six have a native resolution of 20 m x 20 m (B5, B6, B7, B8a, B11 and B12) and the remaining two that we used have a 60 m x 60 m resolution (B1 and B9). We resampled all layers to a common 25 m x 25 m resolution to have the same dimensions as slope and the derived deepSSF layers of NDVI, canopy cover and herbaceous vegetation

**Figure A4:**
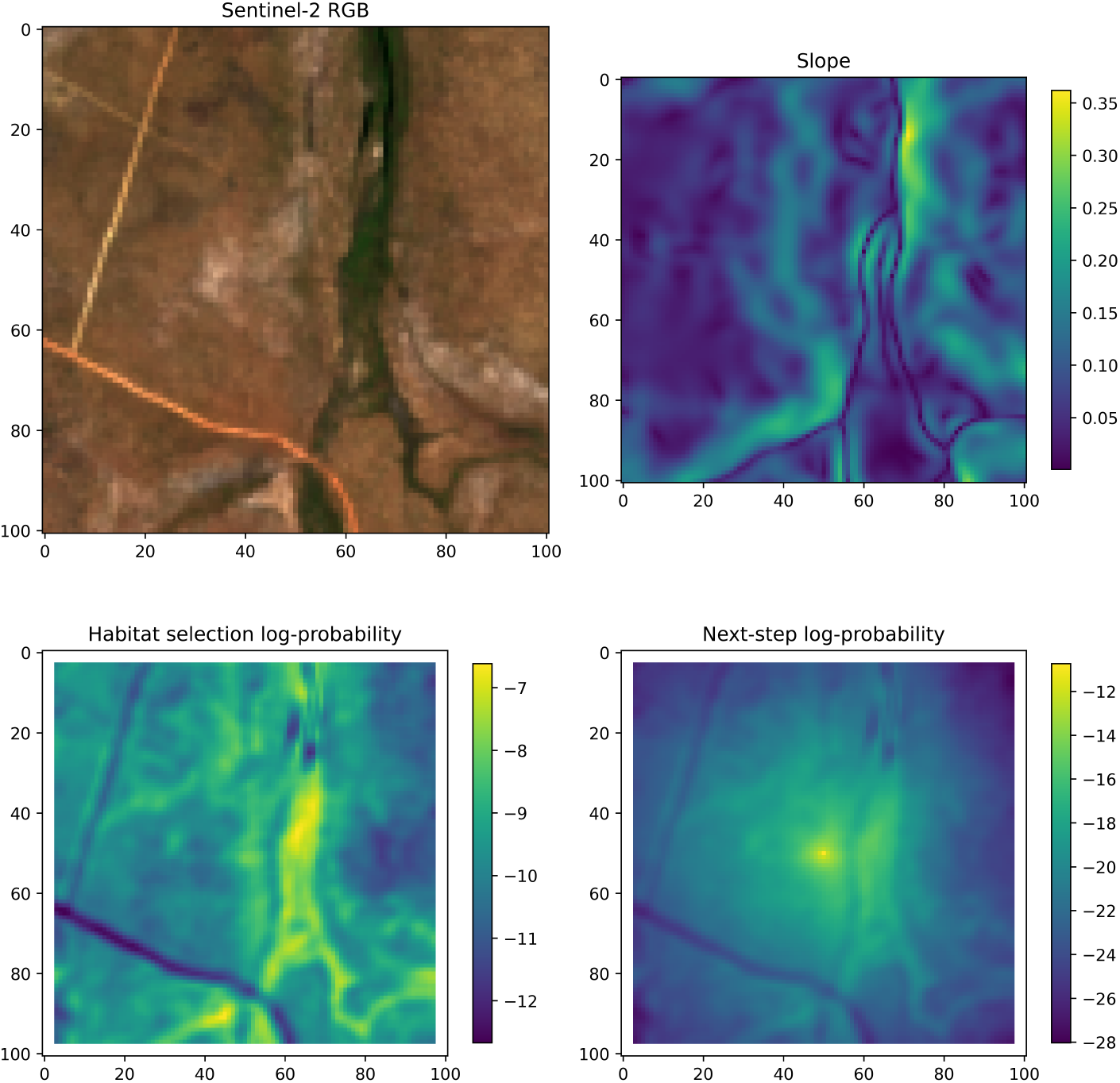
An example of the predicted habitat selection and next-step probability surfaces for 3pm on Day 284 of 2018 (October 11th 2018), which is during the late dry season 2018. The model shows selection for a vegetated area in a valley, and avoidance of roads.

**Figure A5:**
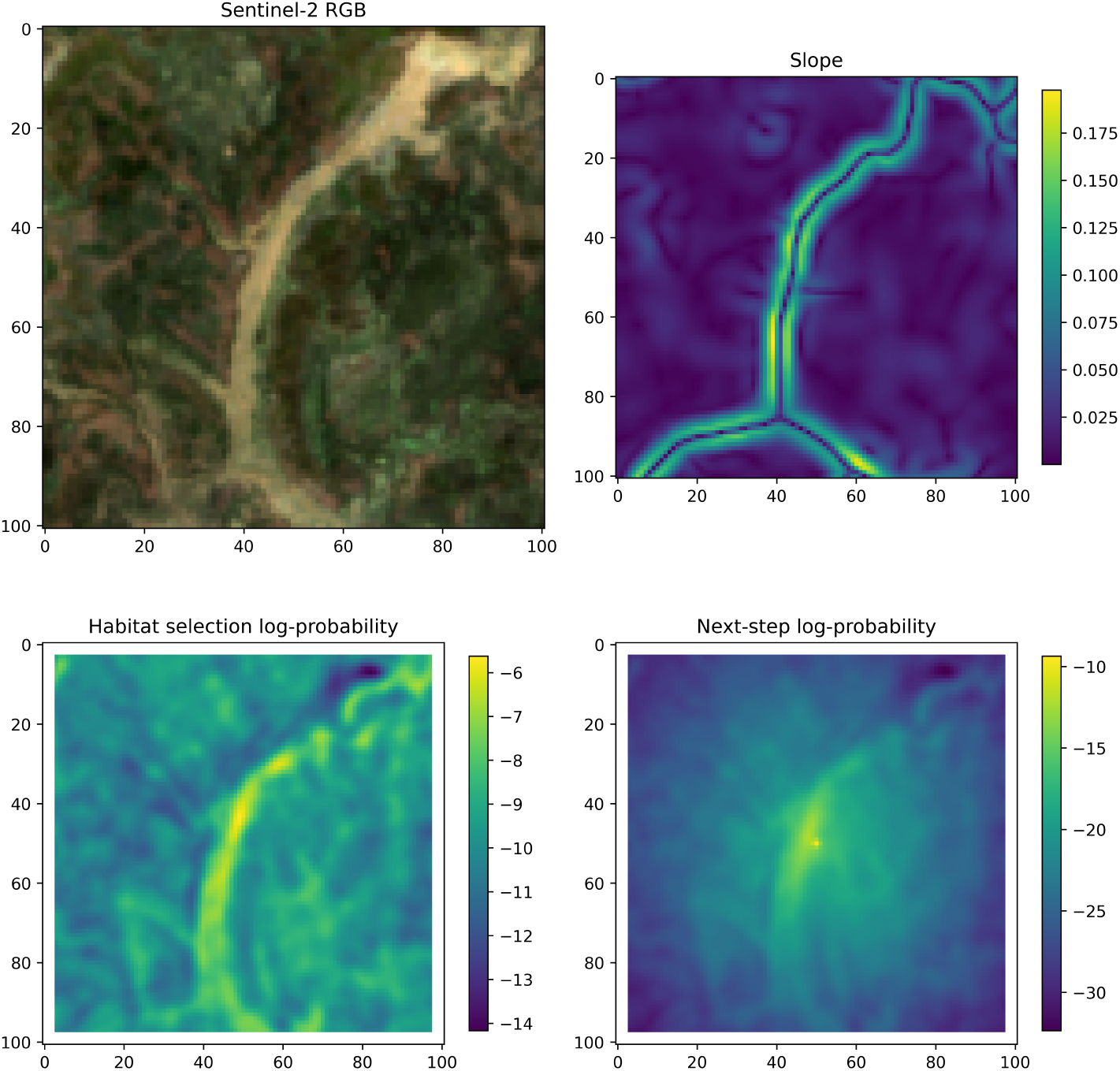
An example of the predicted habitat selection and next-step probability surfaces for 11am on Day 89 of 2019 (March 30th 2019), which is during the late wet season 2019. The model shows selection for a flooded river.

**Figure A6:**
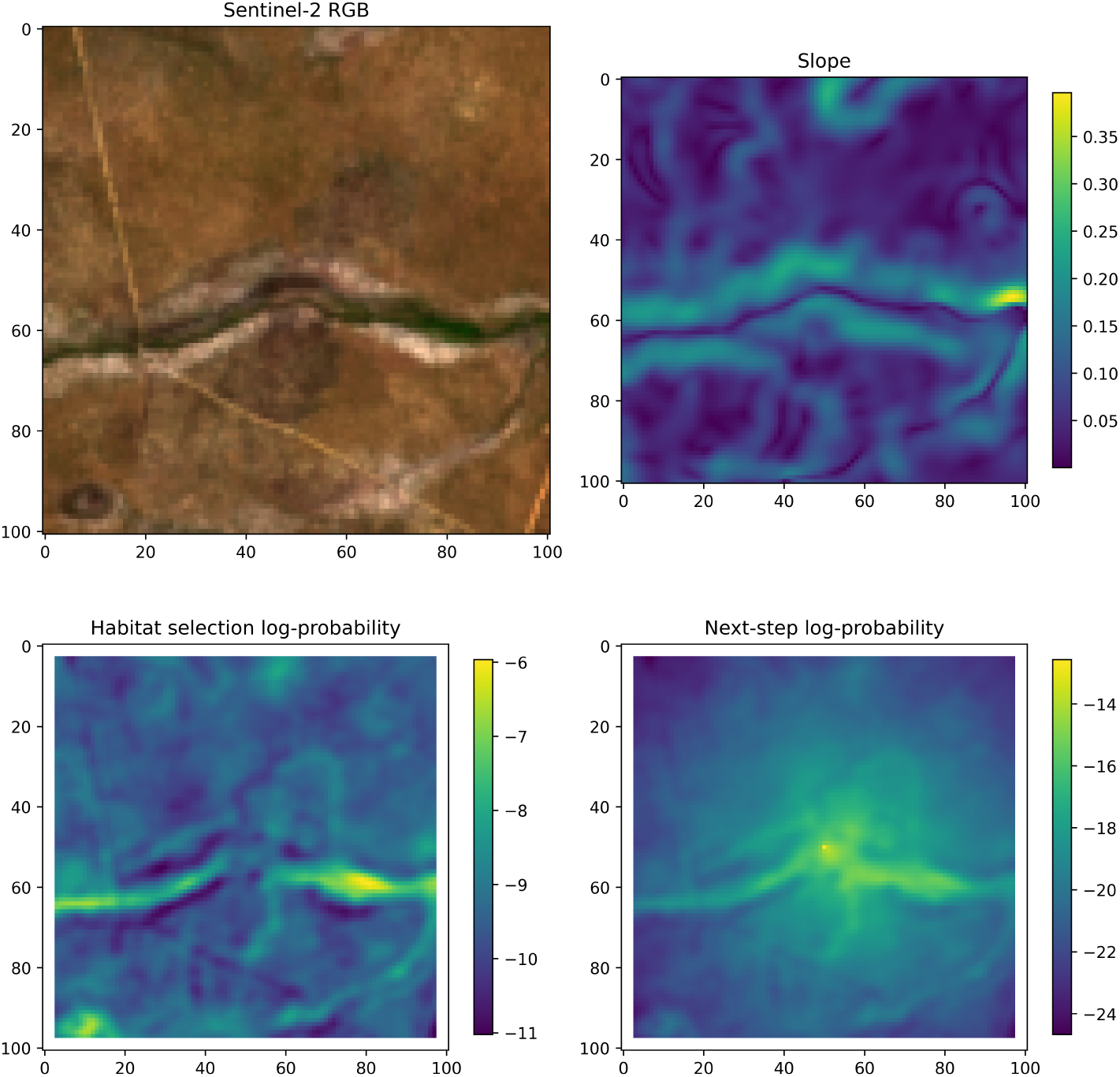
An example of the predicted habitat selection and next-step probability surfaces for 8am on Day 301 of 2019 (October 28th 2019), which is during the late dry season 2019. The model shows selection for a vegetated area in a valley, and a waterhole in the lower left.

**Figure A7:**
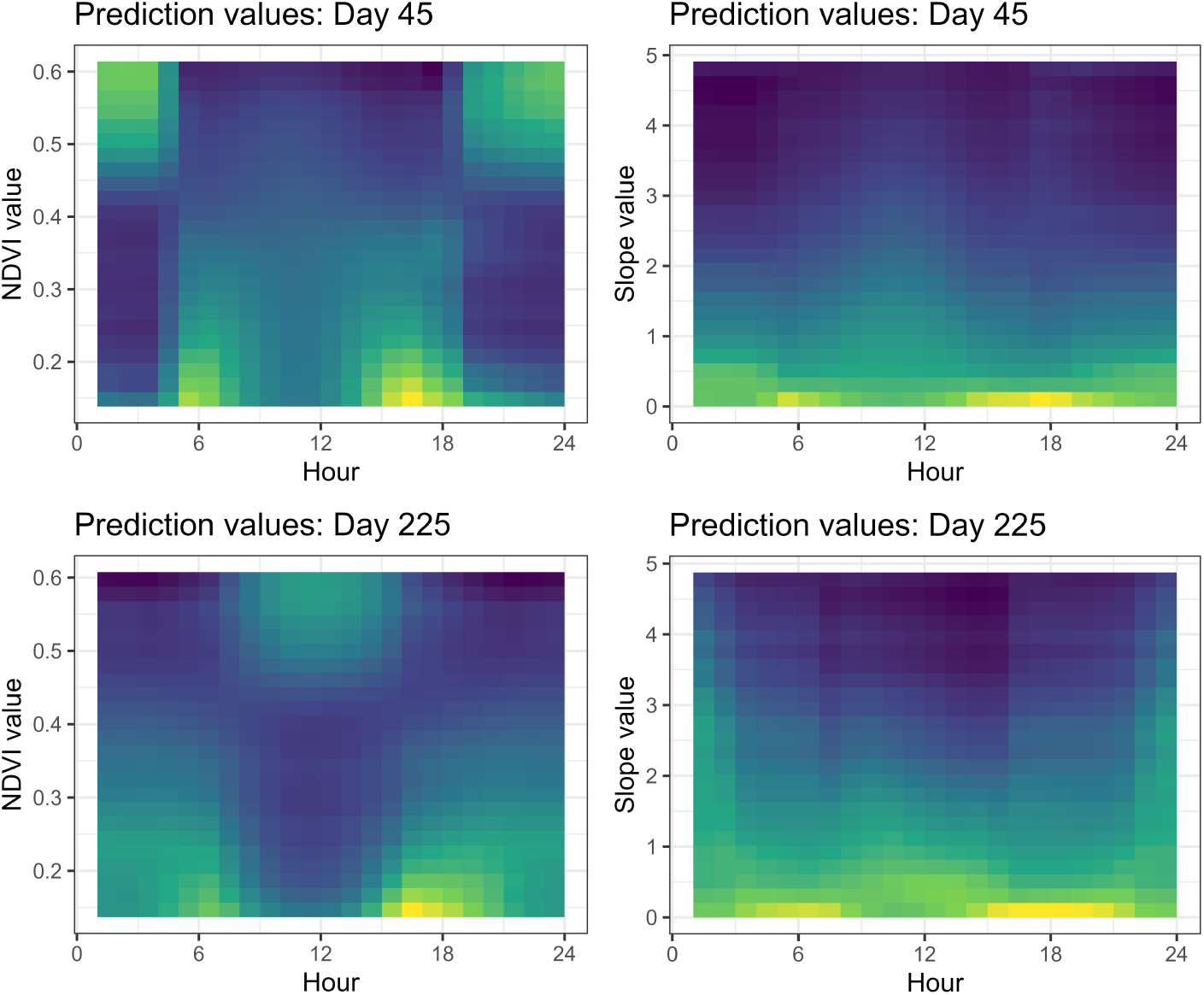
To visualise the effect of particular covariates on the habitat selection probabilities, and how that changes throughout the day and between seasons, we plotted the mean habitat selection probability values from the map extent in Figure 6 for bins of the covariate values. This approach approximates their marginal contribution to the next-step probabilities. We truncated the covariate values to between the 0.01 and 0.99 quantiles to avoid erroneous interpretation from very few values in the tails. We chose Day 45 (February 14th, 2019) and Day 225 (August 13th, 2018) as they are in the middle of the wet and dry seasons, respectively. For Normalised Difference Vegetation Index (NDVI), the pattern of selection around dawn and dusk is similar, although in the wet season there is selection for high values of NDVI during the night, and low values during the middle of the day, which is the opposite in the dry season. For slope, there is consistent selection against high values, which appears stronger in the wet season, and there is selection for very flat areas during the high movement periods at dawn and dusk.

**Figure A8:**
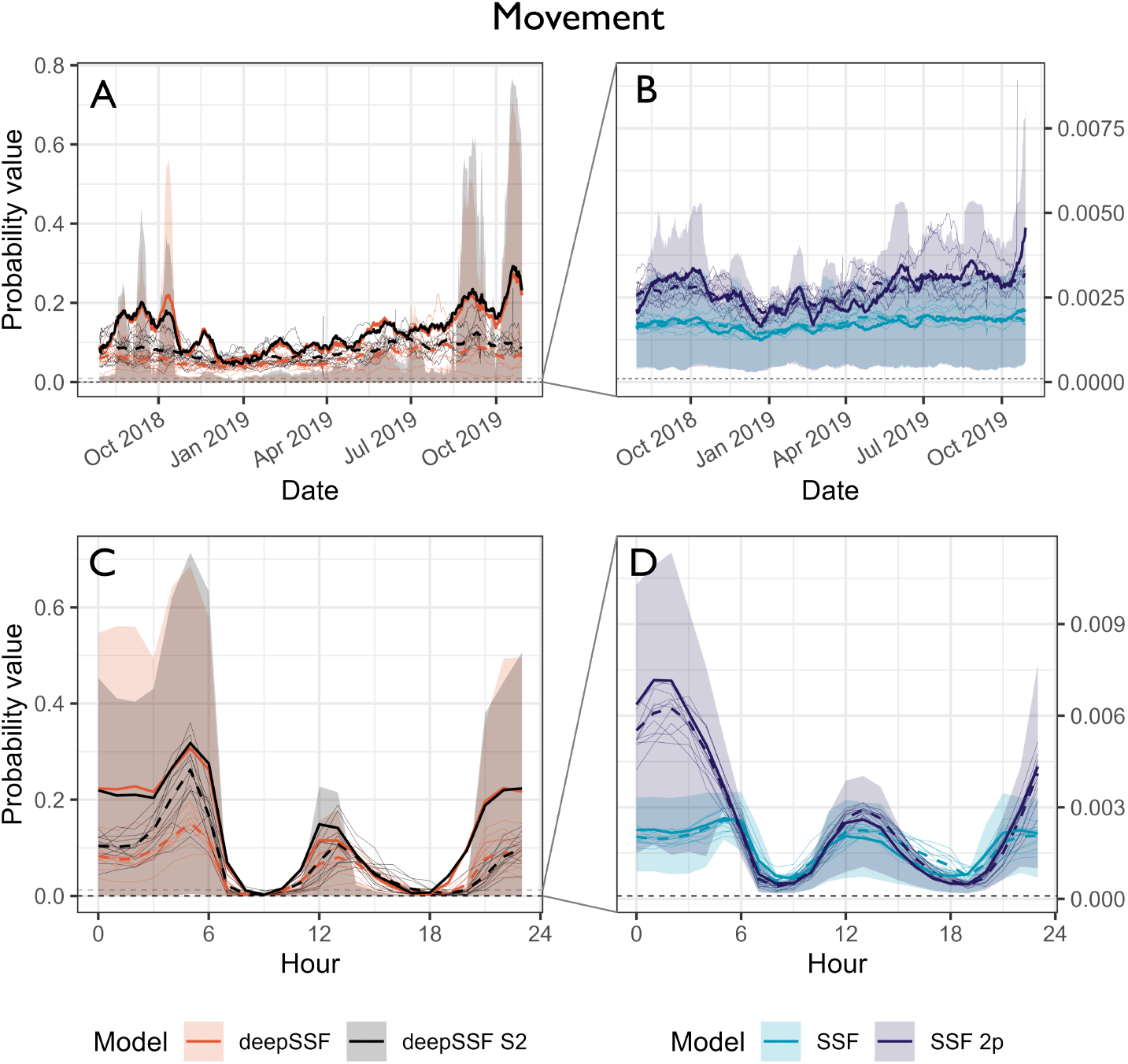
Probability values for the normalised movement process at the observed location of the next-step. We compare between the deepSSF model with derived covariates (Normalised Difference Vegetation Index (NDVI), canopy cover, herbaceous vegetation and slope), the deepSSF model with slope and 12 Sentinel-2 bands at 25 m x 25 m cell resolution (deepSSF S2), a typical SSF fitted to the derived covariates, and an SSF with the derived covariates and temporal dynamics over a daily period using two pairs of harmonics, as presented by Forrest et al. (2024). Panels **A** and **B** show the normalised movement probabilities across the duration of the tracking period, smoothed using a moving window that was 15 days in duration and incremented by one day, and panels **C** and **D** show the normalised movement probabilities across the hours of the day, where the probability values were binned into hour. The solid coloured lines show the average probability for the focal individual that the model was fitted to, and the shaded ribbon is the 50% quantile (there is high variability between probability values, so for clarity we omitted the 95% quantiles). The thin coloured lines are the average probability values for 12 individuals that the model was not fitted to, and are therefore out-of-sample validation data. The dashed coloured lines are the mean values for each model for all of the out-of-sample individuals. The SSF probability values were much lower than the deepSSF values (in both the positive and the negative direction), and the SSF plots in panels **B** and **D** are therefore zoomed in, as indicated by the dashed grey lines in panels **A** and **C**. The accuracy of the movement predictions are fairly consistent throughout the year for both the deepSSF and SSF models, although there is significant variation throughout the day. All models are most accurate outside of the dawn and dusk periods, which are when buffalo have much larger average steps. The larger step length distributions at this time are therefore spread over many more cells, resulting in lower absolute probabilities. The deepSSF models have orders of magnitude larger predicted probability values, likely due to the mixture distribution allowing for a step length distribution with most of the probability mass in just a few cells near zero distance, which is when the buffalo is mostly stationary.

**Figure A9:**
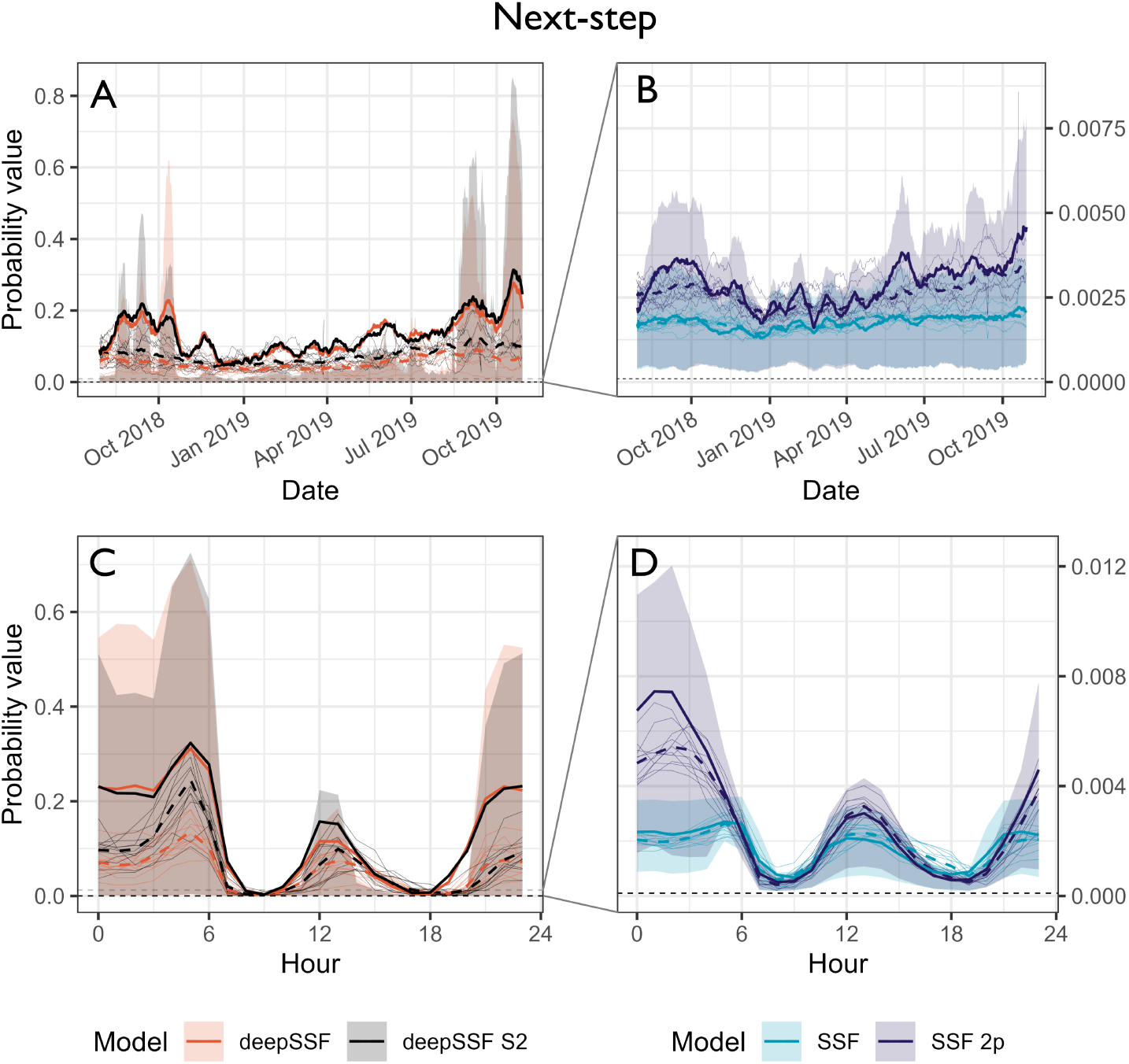
Probability values for the normalised habitat selection process at the observed location of the next-step. We compare between the deepSSF model with derived covariates (Normalised Difference Vegetation Index (NDVI), canopy cover, herbaceous vegetation and slope), the deepSSF model with slope and 12 Sentinel-2 bands at 25 m x 25 m cell resolution (deepSSF S2), a typical SSF fitted to the derived covariates, and an SSF with the derived covariates and temporal dynamics over a daily period using two pairs of harmonics, as presented by Forrest et al. (2024). Panels **A** and **B** show the normalised habitat selection probabilities across the duration of the tracking period, smoothed using a moving window that was 15 days in duration and incremented by one day, and panels **C** and **D** show the normalised habitat selection probabilities across the hours of the day, where the probability values were binned into hour. The solid coloured lines show the average probability for the focal individual that the model was fitted to, and the shaded ribbon is the 50% quantile (there is high variability between probability values, so for clarity we omitted the 95% quantiles). The thin coloured lines are the average probability values for 12 individuals that the model was not fitted to, and are therefore out-of-sample validation data. The dashed coloured lines are the mean values for each model for all of the out-of-sample individuals. The SSF probability values were much lower than the deepSSF values (in both the positive and the negative direction), and the SSF plots in panels **B** and **D** are therefore zoomed in, as indicated by the dashed grey lines in panels **A** and **C**. Due to the large probability values given by the movement process, the next-step probability values are mostly comprised of that contribution, and the inference is similar to that for the movement predictions.

## Notes

https://swforrest.github.io/deepSSF/

### Competing Interest Statement

The authors have declared no competing interest.

